# An evolutionarily conserved odontode gene regulatory network underlies head armor formation in suckermouth armored catfish

**DOI:** 10.1101/2021.06.21.449322

**Authors:** Shunsuke Mori, Tetsuya Nakamura

## Abstract

Odontodes, i.e., teeth and tooth-like structures, consist of a pulp cavity and dentine covered by a mineralized cap. These structures first appeared on the outer surface of vertebrate ancestors and were repeatedly lost and gained across vertebrate clades; yet, the underlying genetic mechanisms and trajectories of this recurrent evolution remain long-standing mysteries. Here, we established suckermouth armored catfish (*Ancistrus sp.*; Loricariidae), which have uniquely evolved dermal odontodes (dermal denticles) all over most of their body surface, as an experimental model animal amenable to genetic manipulation for studying odontode development. Our histological analysis showed that suckermouth armored catfish develop dermal denticles through the previously defined odontode developmental stages. *De novo* transcriptomic profiling identified the conserved odontode genetic regulatory network (oGRN) as well as unique expression of *paired like homeodomain 2* (*pitx2*), previously characterized as an early regulator of oGRN in teeth, in developing dermal denticles. Knockdown of *pitx2* perturbed formation of the epithelial placode of dermal denticles and altered expression oGRN genes. By comprehensively identifying the genetic program for dermal odontode development in suckermouth armored catfish, this work illuminates how dermal odontodes independently evolved and diverged in distinct teleost lineages.

**Summary statement:** Cranial dermal denticles in suckermouth armored catfish develop via an evolutionarily conserved and unique odontode genetic regulatory network.

## INTRODUCTION

Exoskeletal armor is a prime protective adaptation for thriving in a natural environment, and dermal odontodes are the most common exoskeletal feature found across the vertebrate phylogeny of extinct and extant fish. Dermal odontodes are constructed from various types of dentine sometimes covered by enamel/enameloid and primarily function as protectants and ornaments on the vertebrate body (Janvier, 1996). They form body armor in concert with underlying dermal bony plates in odontode-bearing fish, excepting modern cartilaginous fish (Miyake et al., 1999).

During vertebrate evolution, the first dermal odontodes shielding the body surface appeared in in Silurian (∼460 million years ago) in pteraspidomorphi, extinct jawless fish group including anaspids or heterostracans (Keating and Donoghue, 2016). Thereafter, throughout the evolution of jawless and jawed fish, the architecture of odontodes with underlying dermal bony plates has been continuously modified and displays extraordinary morphological divergence (Keating et al., 2015). Odontodes may have been recruited into the oral cavity and transformed into teeth in the “outside-in” model during the origin of jawed fish (Reif, 1982a). However, due to gaps in the fossil record, the evolutionary origin of teeth continues to be disputed and revisited (Berio and Debiais-Thibaud, 2021; Fraser et al., 2010; Witten et al., 2014).

Extant chondrichthyans (cartilaginous fish) and some species of primitive actinopterygians (ray-finned fishes, e.g. gars and bichirs) and sarcopterygians (lobe-finned fish, e.g., coelacanth) retain dermal odontodes on the body surface in the forms of placoid scales or ganoid scales (Huysseune and Sire, 1998; Reif, 1982b; Williamson, 1849; Williamson, 1851). However, dermal odontodes were lost in the early divergence of teleosts (the largest infraclass in actinopterygians) and sarcopterygians; thus, most of them no longer have these structures. Intriguingly, dermal odontodes independently re-emerged as convergent evolution in four living teleost groups: *Xiphioidae* (swordfish), *Denticeps* (denticle herring), *Atherion* (pickleface hardyhead), and *Loricarioidae* (armored catfish) (Fierstine, 1990; Sire and Allizard, 2001; Sire and Huysseune, 1996; Sire et al., 1998). In these fishes, dermal odontodes on the extraoral skin display the conserved odonotode organization and components, e.g., a pulp cavity surrounded by a dentin cone but without an enamel cap. The exceptional resemblance of dermal odontode architectures among these fishes implies a redeployment of the genetic network for odontode formation from existing teeth or resurrection of the genetic program for lost dermal odontodes in each lineage, yet little is known about the genetic basis underlying odontode convergent evolution in teleosts.

The developmental process of odontodes and its core set of regulatory genes were uncovered principally by studying mammalian tooth formation. Tooth development begins with epithelial placode formation; the thickened epithelium with reduced cell proliferation functions as a signaling center, expressing the transcription factor *paired like homeodomain 2* (*pitx2*) as well as genes encoding secreted factors, e.g., *sonic hedgehog* (*shh*), *fibroblast growth factors* (*fgfs*), *bone morphogenic proteins* (*bmps*), and *wnt* (Bei et al., 2000; Bitgood and McMahon, 1995; Chen et al., 2009; Dassule and McMahon, 1998). Via these diffusible molecules, the placode triggers the sequential and reciprocal epithelial-mesenchymal interactions and regulates formation of the tooth primordium, i.e., tooth germ, through the well-defined developmental stages of bud, cap, bell, and eruption (Balic and Thesleff, 2015; Thesleff and Tummers, 2008).

Recent comparative studies established histological and genetic links between teeth and various dermal odontodes. For example, placoid scales, the dermal odontodes of cartilaginous fish, form through the strictly conserved bud, cap, bell, and eruption stages (Debiais-Thibaud et al., 2015). Also, *shh*, *wnt*, *bmp*, *fgf* and other genes indispensable for tooth development are expressed in a comparable manner during both tooth and placoid scale formation (Cooper et al., 2017; Cooper et al., 2018; Debiais-Thibaud et al., 2015; Debiais-Thibaud et al., 2011). These architectural and molecular correspondences between tooth and dermal odontode development supports the classical hypothesis that genetic cascades were co-opted from dermal odontodes to teeth (or vice versa). However, due to the lack of experimental model animals amenable to genetic manipulation, there is little direct experimental evidence for this hypothesis and, more broadly, for how odontodes are recruited into novel tissues.

The catfish order (Siluriformes) offers a remarkable and unique opportunity to understand the evolutionary mechanisms of dermal odontodes. The last common ancestor of catfish is thought to have lost all body scales, but some lineages in the superfamily Loricarioidea independently reverted to gain dermal odontodes, i.e., dermal denticles, with/without underlying dermal bony plates (Haspel et al., 2012; Sire and Huysseune, 1996). Although these dermal denticles in catfishes originated via convergent evolution, their structures display the rigorously conserved odontode characteristics albeit lacking surface enamel. The distribution of dermal denticles along the body in these catfish lineages is evolutionarily diverse, occuring on the head, trunk, or both (Rivera-Rivera and Montoya-Burgos, 2017). These disntinct lineages with their independent and morphologically-diverse gains of dermal denticles are prominent model systems for studying the genetic mechanisms of dermal odontode evolution. Furthermore, the genomic sequences of three catfish species (*Ictalurus punctatus, Pterygoplichthys pardalis, Platydoras armatulus*) with/without dermal denticles were recently assembled, establishing the catfish order as a genetically approachable clade (Liu et al., 2016). This combination of unique, yet anatomically conserved, odontode development and available genomic resources make the catfish order ideal for identifying the genetic mechanisms of dermal odontode evolution.

In this study, we established the suckermouth armored catfish (*Ancistrus sp.*; Loicariidae) as an experimental model animal to investigate the evolutionary and developmental mechanisms of dermal odontodes. Our histological studies revealed that cranial dermal denticles continuously arise and develop from embryonic stages to, at least, the early adult stage via the previously defined developmental stages of odontodes. We then compared gene expression profiles of cranial, scute, and oral tissues by *de novo* transcriptomic assembly and identified both evolutionarily conserved and unique elements of a genetic regulatory network for dermal denticle formation. Finally, we demonstrated that functional knockdown of *paired like homeodomain 2* (*pitx2*), uniquely expressed in the epithelial placode of dermal denticles, reduced cranial denticle number and size as well as the expression of genes involved in the odontode genetic regulatory network (oGRN). These results highlight the conserved and unique genetic mechanisms of dermal denticle formation in *Ancistrus sp.* and shed light on how dermal denticles have repeatedly evolved along the vertebrate body.

## RESULTS

### Cranial dermal denticle development in *Ancistrus sp*

Suckermouth armored catfish are unique among catfish in that they have evolved dermal denticles over most of their bodies except for the abdomen (Alexander, 1966). Previous studies showed that dermal denticles in suckermouth armored catfish possess conserved odontode architecture (Rivera-Rivera and Montoya-Burgos, 2017). To examine the process of dermal denticle development, we performed mineralization staining and histological analysis of developing dermal denticles in *Ancistrus sp*., commonly called the bristlenose pleco and known for its relatively small size and ease of breeding. Cranial dermal denticles were the first denticles to appear and formed on the cranial skin at 96 hours post fertilization (hpf) (Fig. 1A) as revealed by alkaline phosphatase (ALP) staining, a marker of odontoblast differentiation (Magne et al., 2004). We observed eight symmetrically arranged dermal denticles on both the left and right sides of the cranium (Fig. 1A). These cranial denticles were not stained by alizarin red, reflecting their pre-mineralized state (Fig. S2A). At this stage, based on hematoxylin and eosin (HE) staining, the cranial dermal denticles are anchored to underlying cartilages via attachment-fiber like structures, as previously described for the placoid scales of cartilaginous fish (Fig. 1B) (Miyake et al., 1999) and were shed approximately by 10-14 days post fertilization (dpf) (the early larva stage) (data not shown), thus we term them the primary dermal denticles (Fig. 1C). After 14 dpf when the larvae have consumed almost all yolk contents and begun feeding, secondary cranial dermal denticles appear at new positions on the cranial skin surface (Fig. S2B). These denticles as well as the underlying skull roof displayed mineralization based on Alcian blue and Alizarin red staining (Fig. 1D). As the juveniles grew, the number of mineralized secondary cranial dermal denticles increased along with the expansion of underlying dermal bony plates to cooperatively construct the head armor—the most distinctive feature of armored catfish (Fig. 1D and Figs. S1, S2B, C). Also, the secondary cranial dermal denticles continuously appeared and matured from ∼ 14 dpf (the larval stage) to, at least, 360–540 dpf (the early adult stage, 6–7 cm total length (TL)). In summary, *Ancistrus sp*. develops deciduous primary denticles and consecutive secondary denticles at different developmental time points, and only the latter mineralizes and forms the head armor.

**Fig. 1.**
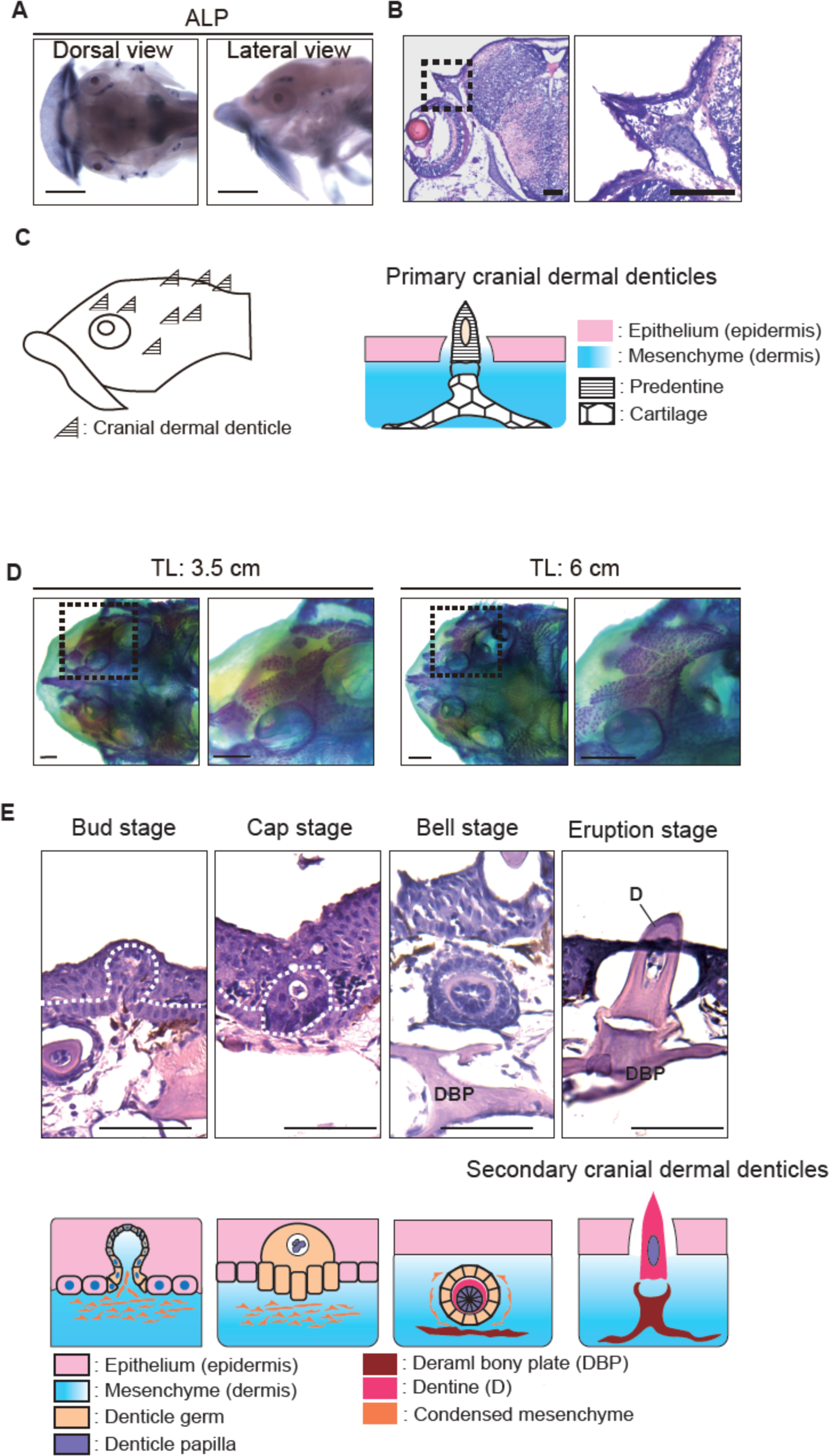
Cranial dermal denticles develop through conserved odontode developmental stages. (A) ALP staining shows developing primary cranial dermal denticles. Eight cranial dermal denticles form on both the left and right sides. Scale bars: 0.5 mm. (B) HE-stained transverse section of *Ancistrus sp*. embryo at 96 hpf. Primary cranial dermal denticles form in the cranial epithelium and mesenchyme, being anchored to underling cartilages via the attachment fiber-like structures. The right panel shows higher magnification of the dotted box area in the left panel. Scale bars: 0.1 mm. (C) Schematic illustration of primary cranial dermal denticle positions (left) and their structure (right). (D) Bone and cartilage staining of a juvenile (TL: 3.5 cm) and adult fish (TL: 6 cm). The number of cranial dermal denticles increased with underlying dermal plate expansion from the juvenile to the adult stage. Each right panel shows higher magnification of the dotted box area in the left panels. Scale bars: 1 mm. (E) Secondary cranial dermal denticle development through the bud, cap, bell, and eruption stages in adult fish (TL: 6 cm, top) and the diagram of the developmental stages (bottom). White dotted lines indicate the boundary between the surface epithelium and cranial dermal denticle germs. Scale bars: 100 µm. At least two biological replicates were investigated in each experiment.

### The developmental stages of cranial dermal denticles in suckermouth armored catfish

To characterize the developmental stages of the primary and secondary cranial dermal denticles, we examined the histology of dermal denticles at 60 hpf (primary denticles at the embryonic stage), 120 dpf (secondary denticles at the juvenile stage), and 360 – 540 dpf (secondary denticles at the early adult stage). In adult fish, secondary dermal denticles at the bud stage exhibit a thickened basal layer of the epithelium just beginning to invaginate towards the surface epithelium (Fig. 1E). Subsequently, at the cap stage, the epithelium becomes more folded and encloses the denticle papilla, forming a dermal denticle primordia termed the denticle germs (Welten et al., 2015). The epithelial cells then completely encircle the denticle papilla where predentin differentiation initiates (the bell stage) (Fig. 1E). The underlying dermal bony plates simultaneously begin to development in the mesenchymal layer under the denticle germs. Finally, dentin cones erupted from the epithelium, being directly associated with underlying dermal bony plates (the eruption stage) (Fig. 1E). It was difficult to reliably identify all four developmental stages in embryos and juveniles, but the cap stage was detected in embryos and the cap, bell, and eruption stages in juveniles (Fig. S2D). From our histological analysis, we conclude that, in *Ancistrus sp.,* both primary and secondary cranial denticles develop through the conserved odontode developmental stages as previously defined in tooth and placoid scale morphogenesis (Fig. 1E) (Berio and Debiais-Thibaud, 2021; Ellis et al., 2015; Thesleff and Tummers, 2008).

### *De novo* transcriptomic profiling of teeth and cranial and trunk denticles in *Ancistrus sp*

Because the developmental stages of *Ancistrus sp*. cranial dermal denticles are shared with those of teeth and placoid scales of other vertebrates, we decided to test for conservation of the odontode gene regulatory network (oGRN) in these fish (Fraser et al., 2010). To comprehensively identify the genes that function in *Ancistrus sp*. odontode development, we conducted comparative *de novo* RNA sequencing assembly (*de novo* RNA-seq) of cranial dermal denticles (juveniles and adult), trunk dermal denticles on scutes (adult), and teeth (adult). Because the odontodes are small for manual dissection, we recovered total RNA from cranial skin containing dermal denticles at 120 dpf (juveniles, approximately TL 2–3 cm, “CDj”) and at ∼360 dpf (sexually matured adult, approximately TL 5–6 cm, “CD”) as well as adult scutes containing trunk dermal denticles (“Scute”), and adult oral epithelial tissues with teeth (“Mouth”) (Fig. S3A).

After *de novo* assembly of short reads from high-throughput sequencing, the contigs were further processed to longer fragments, i.e., unigenes, which were functionally annotated by the NCBI Nr database. First, we investigated the species distribution of homologues for the annotated unigenes and found that 39.8% of the unigenes were homologues of channel catfish genes (*Ictalurus punctus*) and 17.3% were homologues of red piranha genes (*Pygocentrus nattereri*) (Fig. S3B), suggesting that the genes obtained from *Ancistrus sp. de novo* RNA-seq are more similar to those of the catfish family than to those of any other fish species. All assembled unigenes from the developing dermal odontodes of *Ancistrus sp*. were categorized into three Gene ontology (GO) groups: biological process (BP), cellular component (CC), and molecular function (MF) (Fig. S3C). In the BP category, 7,739 unigenes were assigned to “developmental process” (Fig. S3C), confirming that *de novo* RNA-seq successfully enriched for genes involved in the developmental processes of body structures.

Hierarchical clustering of the top 25,000 differentially expressed unigenes across all samples indicated that the gene expression profile of cranial dermal denticles (CDj and CD) is more similar to that of trunk dermal denticles (Scute) than to that of teeth (Mouth) (Fig. 2A). To explore the similarity of functional gene categories among the CD, CDj, Scute, and Mouth samples, we then carried out GO enrichment analysis, comparing CDj to each adult tissue. The analysis showed that GO:0048856 “the anatomical structure development”, which includes the gene set associated with odontogenesis, was enriched in all adult odontode tissue (Fig. 2B and Fig. S3D, asterisks).

**Fig. 2.**
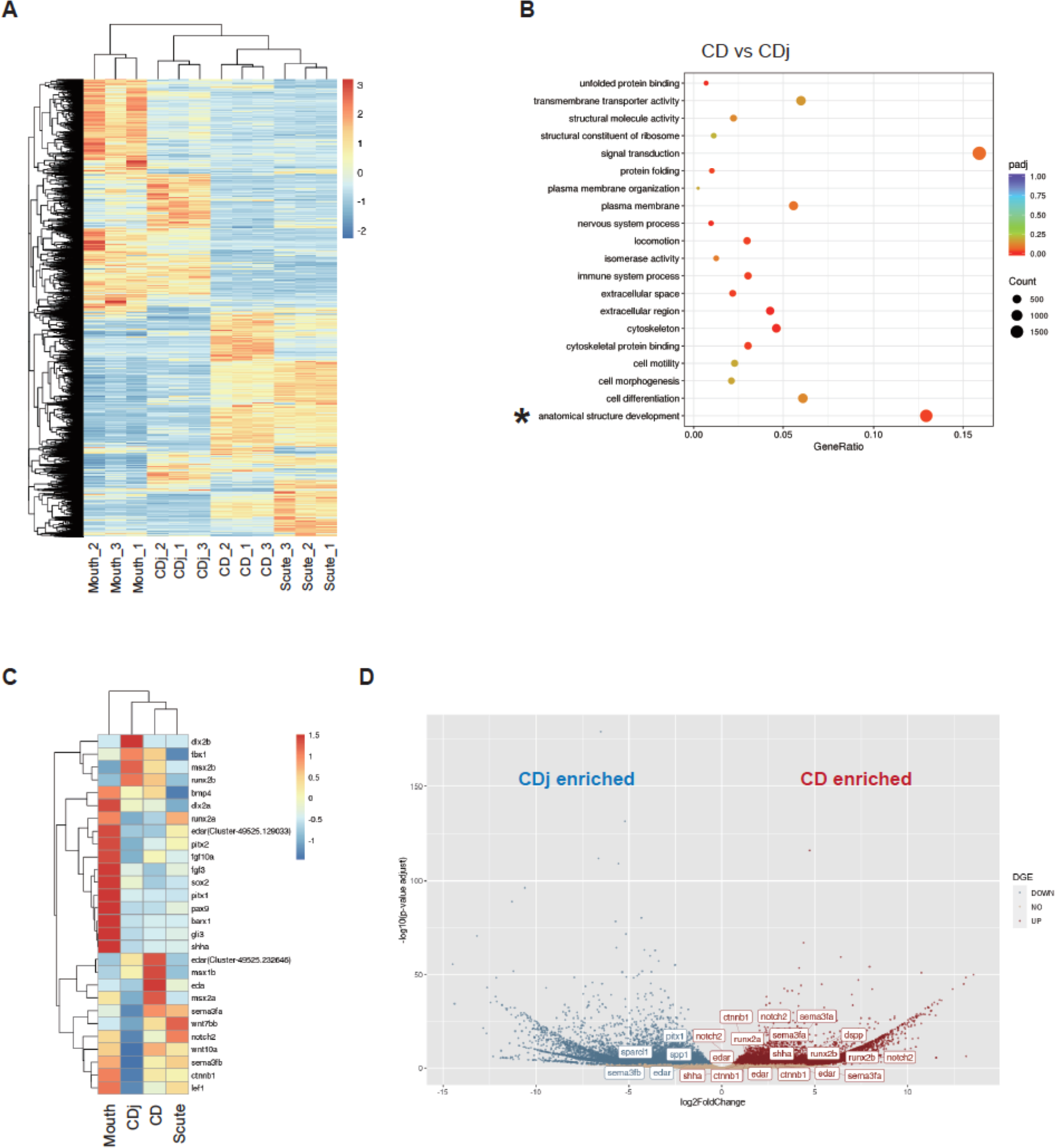
Comprehensive identification of the genes involved in dermal denticle formation of *Ancistrus sp*. Differential gene expression analysis was performed using *de novo* RNA-seq data from juvenile cranial dermal denticles (CDj, *n*=3), adult cranial dermal denticles (CD, *n*=3), Teeth (Mouth, *n*=3), and scutes (Scute, *n*=3). (A) The heatmap of hierarchical clustering of top 25,000 differentially expressed unigenes. Note that all dermal odontode samples (CDj, CD, and Scute) were clustered together with Mouth as an outgroup. (B) GO enrichment analysis of the upregulated genes in CD compared to CDj. The vertical axis is the GO terms, and the horizontal axis is the enrichment of each term. The size of each point represents the number of upregulated genes, and the color of the points represents the adjusted *p*-value. (C) The heatmap produced by hierarchical clustering of the expression levels of 27 selected genes involved in oGRN. The color bars indicate the relative expression levels in FPKM. (D) The volcano plot displays the pattern of gene expression values for CD relative to CDj. Significantly enriched genes in CDj are highlighted in blue (log2foldchange < −0.6 and adjusted *p*-value < 0.01) and enriched genes in CD are in red (log2foldchange > 0.6 and adjusted *p*-value < 0.01). The gene symbols show differentially expressed genes associated with oGRN and odontogenesis. As different unigenes were annotated by the same gene names or even the homologous genes in different species, the multiple same gene names were shown in the plot.

Previous studies indicated that the oGRN is functional in both tooth and dermal denticle development (Fraser et al., 2010). To test whether the oGRN underlies odontode formation in *Ancistrus sp*., we examined the expression level of 27 genes previously reported as the central components of the oGRN in early dermal odontode and tooth formation (Fig. 2C). Hierarchical clustering of all samples with only the representative oGRN genes grouped the CDj, CD, and Scute samples together with Mouth as an outgroup (Fig. 2A, C). As we anticipated, oGRN genes were highly enriched in CDj and CD samples, including *msx1b, msx2a/b, dlx2a/b, bmp4,* and *edar.* Notably, *wnt7bb, wnt10a, ctnnb1* (*β-catenin*), and *lef1*, the components of the Wnt/*β*-catenin signaling pathway indispensable for odontode mineralization, were significantly enriched in adult odontodes (CD, Scute, and Mouth samples) (Fig. 2C) (Chen et al., 2009). Intriguingly, some samples had distinct expression levels of certain oGRN genes, i.e., *bmp4* and *dlx2a* were highly enriched in CDj, CD, and Mouth but not in Scute. Also, the Mouth sample highly expressed the early odonotogenesis regulators (e.g., s*hha* and *pitx2)* and the dental epithelial stem cell marker (*sox*2), a gene indicative of polyphyodonty (teeth replacement throughout life) (Martin et al., 2016). Thus, in suckermouth armored catfish, the conserved oGRN seems to function in early cranial and trunk dermal denticle development as it does during tooth and dermal odontode development in other vertebrates.

Next, to investigate the similarity of the maturation process of teeth and dermal odontodes, we created volcano plots and compared gene expression levels in the CDj, CD, Scute, and Mouth samples (Fig. 2D and Fig. S3E). The expression of *sparc-like 1* (*sparcl1*) and *secreted phosphoprotein 1* (*spp1*), which are indispensable for mineralization of dentine and bones (Enault et al., 2018; Kawasaki et al., 2004; MacDougall, 1998), were higher in CDj than in CD, Scute, and Mouth (Figs. 2D and Fig. S2E). In contrast, *runx2a* (*runt-related transcription factor2a*), *β-catenin* (*ctnnb1*), *edar*, and *notch2*, factors indispensable for odontoblast differentiation (Cai et al., 2011; Chen et al., 2009; Gaur et al., 2005; Li et al., 2011; Tucker et al., 2000), were enriched in CD, Scute, and Mouth. Moreover, *dentin sialophosphoprotein* (*dspp*), which is abundant in the dentin extracellular matrix (MacDougall, 1998; Yamakoshi, 2008), was also more highly expressed in these adult samples than in CDj. The function of these genes in dentine mineralization and maturation requires more comprehensive investigation. Still, our data implies that Sparcl1 and Spp1 contribute to the early mineralization process of dentine, and DSPP, previously reported to be regulated by the Wnt-Runx2 pathway (Gaur et al., 2005; Komori, 2010), controls the late phase of odontoblast differentiation and mineralization. Overall, there is conservation between the genetic programs involved in the maturation and mineralization of dermal denticles in suckermouth armored catfish and those programs regulating tooth and dermal odontode development in other vertebrates (Chen et al., 2005; Enault et al., 2018).

### oGRN genes showed spatially restricted expression patterns in developing cranial dermal denticles of *Ancistrus sp*

We next performed *in situ* hybridization (ISH) with RNA probes complementary to selected oGRN genes (*pitx2, bmp4, dlx2a, fgf3, shha, wnt10a, β-catenin,* and *pax9*) (Fraser et al., 2010). All tested genes were expressed in developing cranial dermal denticles at 96 hpf (primary denticles at the embryonic stage) and at 120 dpf (secondary denticles at the juvenile stage) (Fig. 3A and Figs. S4B, S5) as well as in embryonic teeth and dermal denticles on pectoral fin spines (Fig. S4). In secondary cranial dermal denticles at the eruption stage, the expression of *pitx2, shha,* and *dlx2a* was restricted to the epithelium surrounding dermal germs (Fig. 3A, B). Transcripts for *β-catenin*, *wnt10a, bmp4,* and *fgf3* were detected in both epithelial and mesenchymal cells (Fig. 3A, B). *Pax9* is expressed in the mesenchyme of developing teeth and placoid scales (Fraser et al., 2010), but here was found in both epithelial and mesenchymal cells (Fig. 3A, B). The expression domains of *fgf3* and *pax9* expression domain were shown to be associated with the direction of denticle eruption in shark teeth, and the eruption of cranial dermal denticles may also be regulated by these molecules (Rasch et al., 2016). These results suggested that, despite minor variants, the oGRN genes controlling dermal odontode development in suckermouth armored catfish exhibit patterns comparable to those previously reported for other vertebrate odontodes (Berio and Debiais-Thibaud, 2021; Fraser et al., 2010).

**Fig. 3.**
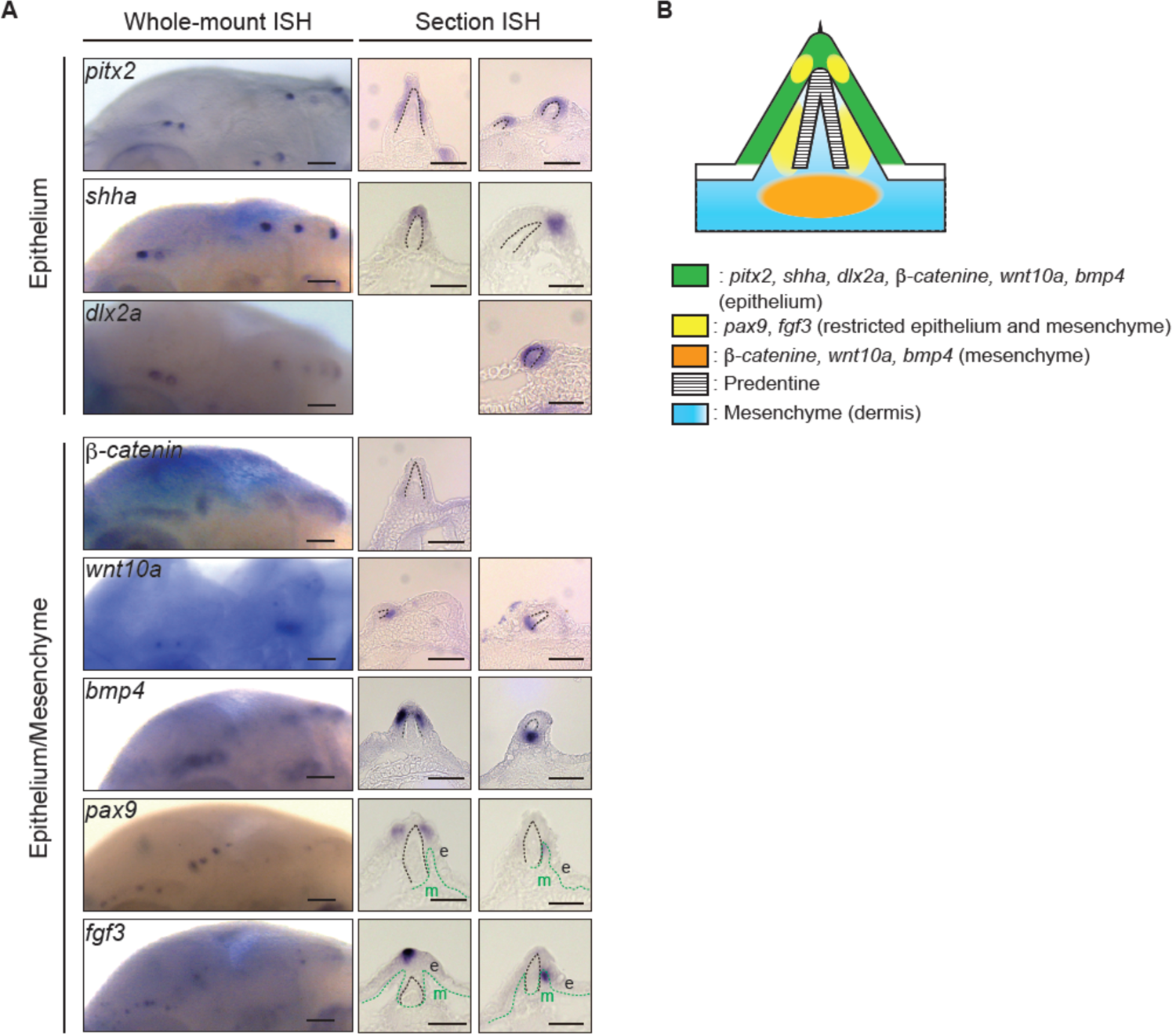
oGRN gene expression in primary cranial dermal denticles. (A) *In situ* hybridization (ISH) for oGRN genes in secondary cranial dermal denticles of *Ancistrus sp*. at 96 hpf. Whole-mount (left) and section (right) *in situ* hybridization of represented oGRN genes at the eruption stage. *Pitx2, dlx2a* and *shha* transcripts were detected in the epithelium. *b-catenin, wnt10a, bmp4, pax9,* and *fgf3* transcripts were in both the epithelium and mesenchyme. Black dotted lines indicate the borderline of the epithelium and dentin cone. Green dotted lines indicate the borderline of the epithelial or dentin cone and mesenchyme. For the epithelium genes, the left and right section panels show the entire epithelial and distal tip expression, respectively. In the epithelium/mesenchyme genes, the left and right section panels show the epithelial with/without mesenchymal expression and mesenchymal gene expression. Note that, to display the representative oGRN gene expression patterns, we selected different slice positions of dermal denticles for some of the section ISH photos. Accordingly, some denticle germ morphology appear to be different from others. Scale bars: 100 µm (left) and 50 µm (right). e; epithelium, m; mesenchyme. (B) Schematic illustration of oGRN gene expression patterns in secondary cranial dermal denticles at the eruption stage. At least three biological replicates were investigated in each experiment.

### Cranial dermal denticles form with the epithelial placode

Our *de novo* RNA-seq and ISH identified *pitx2, dlx2a*, and *bmp4* in the epithelium of dermal germs at the eruption stages in *Ancistrus sp*. (Figs. 2, 3 and Figs. S4, S5). Expression of these genes in the dental placode contribute to the initiation of tooth germ development by inducing the epithelial thickening (Green et al., 2001; St Amand et al., 2000). Thus, we wondered whether a similar developmental program arises within the epithelial placode during dermal denticle development in suckermouth armored catfish. To dissect the induction mechanisms of the epithelial placode for dermal denticles, we examined the expression patterns of *pitx2*, *dlx2a*, *bmp4, pax9,* and *shha* from 48 to 72 hpf when primary dermal denticles begin to form. At 60 hpf, among these genes, only *pitx2* expression was detected in the developing primary cranial dermal denticles (Fig. 4A and Fig. S6A, B) and by 72 hpf, expression of *dlx2a* and *bmp4* were also present (Fig. S6C). From 48-72 hpf, we could not detect the denticle expression of other oGRN genes. e.g., *pax9* or *shha,* in denticles. These genes are reported to be expressed slightly later than *pitx2* in mammalian tooth development (Fig. S6C) (Matalová et al., 2015). We observed a similar stepwise expression pattern of *pitx2, dlx2a,* and *bmp4* in developing teeth and dermal denticles on pectoral fin spines (Fig. S6A–C).

**Fig. 4.**
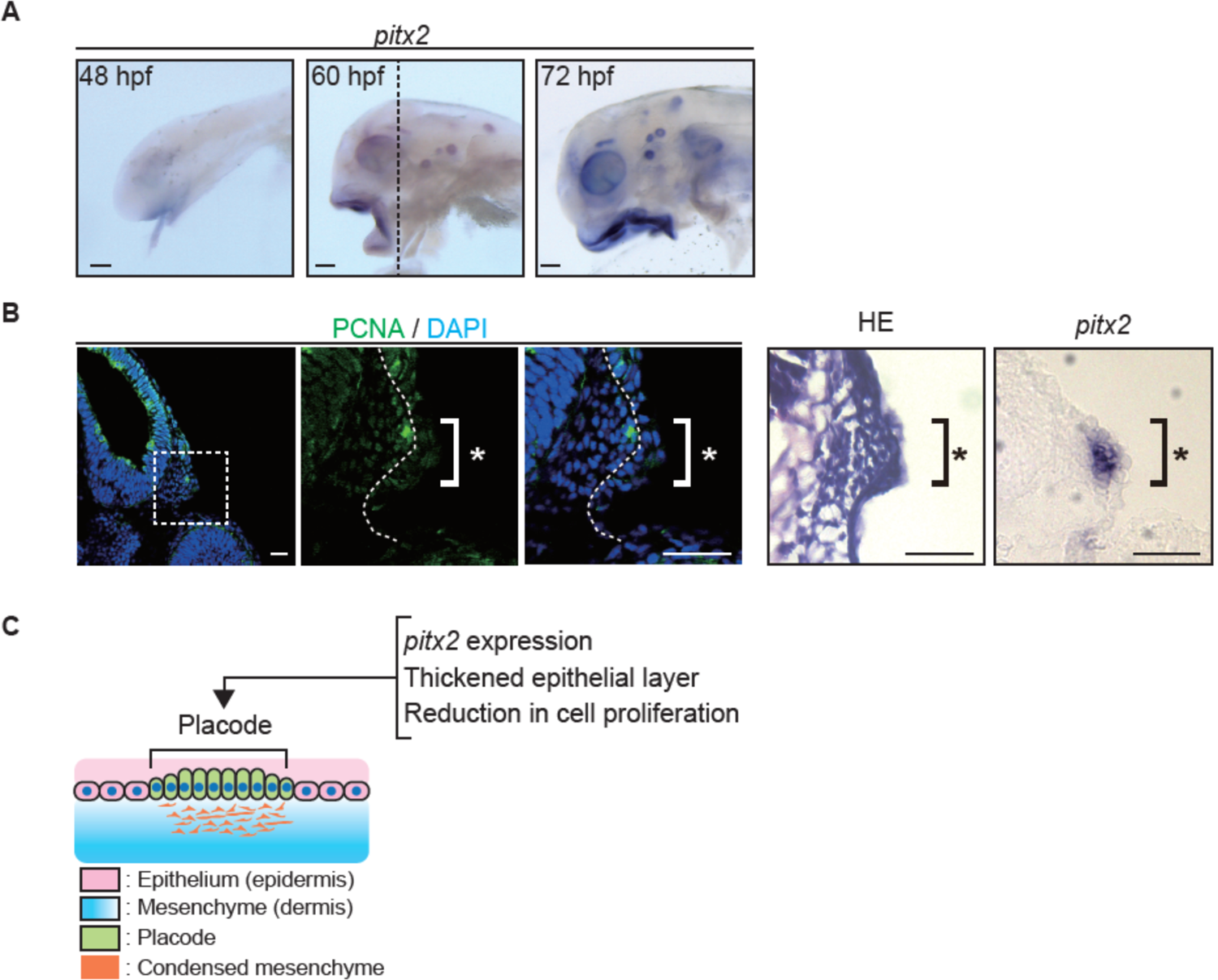
Anatomically and molecularly defined epithelial placode for cranial dermal denticles. (A) Whole-mount *in situ* hybridization of *pitx2* in *Ancistrus sp*. embryos at 48, 60, and 72 hpf. *Pitx2* expression in developing dermal denticles was discerned at 60 dpf. Scale bars: 100 µm. (B) Immunostaining of PCNA with nuclear staining (DAPI), HE staining, and *in situ* hybridization of *pitx2* for transverse sections at the position of the dotted line in A. *Pitx2* transcripts were detected in the anatomically defined placode (asterisk; thickened epithelial layer with the reduction of cell proliferation). Scale bars: 50 µm. (C) Schematic illustration of the cranial dermal denticle placode. At least three biological replicates were investigated in each experiment.

To further characterized the placode formation during cranial dermal denticle development, we performed immunostaining for proliferating cell nuclear antigen (PCNA). A reduction in this stain marks the initial placode formation (Cooper et al., 2017; Di-Poï and Milinkovitch, 2016). PCNA immunostaining revealed reduced cell proliferation in the thickened epithelium relative to the adjacent tissue at 60 dpf (Fig. 4B and Fig. S7). Cryosectioning of embryos used for *pitx2* ISH embryos revealed that *pitx2* was also enriched in the thickened epithelium (Fig. 4B and Fig. S7). These results demonstrated that the anatomically and molecularly defined placodes underlie cranial dermal denticle formation in *Ancistrus sp*. (Fig. 4C).

### Pitx2 establishes the epithelial placode of cranial dermal denticles

Based on our ISH results, *pitx2* is the earliest gene to be expressed in the epithelium of *Ancistrus sp*. dermal denticles. Pitx2 is an inducer of the mammalian tooth epithelial placode (St Amand et al., 2000), thus, could also establish placode formation by inducing oGRN gene expression in *Ancistrus sp*. Given that *pitx2* is required for the development of teeth, eyes, pituitary glands, muscles, lung, and heart in vertebrates, whole-body genetic deletion of *pitx2* could result in embryonic lethality before dermal denticles develop (Lin et al., 1999). To circumvent this possibility, we targeted the *pitx2* mRNA using CRISPR-Cas13d, which was recently reported as a novel gene knockdown tool for targeting maternal and zygotic mRNAs (Kushawah et al., 2020). We injected Cas13d protein with/without guide RNAs complementary to *pitx2* mRNA into one- or two-cell stage embryos (Fig. 5A). By 72 hpf, the level of *pitx2* mRNA in knockdown embryos was 50% of that of control embryos as confirmed by real-time PCR (Fig. 5B). Knockdown embryos exhibited smaller and thinner placodes than control embryos (Fig. 5C), and, based on ISH, these thinned placodes showed a marked decrease in *pitx2* transcripts, histologically validating the real-time PCR result (Fig. 5C). At 96 hpf, both the staining intensity of the odontoblast differentiation marker ALP and the number of dermal denticles were decreased in knockdown embryos (Fig. 5D). Finally, to test whether *pitx2* triggers oGRN gene expression, we investigated the transcript levels of *dlx2a* and *bmp4* and found that *pitx2* knockdown embryos exhibited reduced expression of these genes in cranial dermal denticles, teeth, and denticles on pectoral fin spines (Fig. 5E and Fig. S8). These results demonstrate that Pitx2 induces oGRN gene expression and is responsible for forming the placode of dermal odontodes in suckermouth armored catfish.

**Fig. 5.**
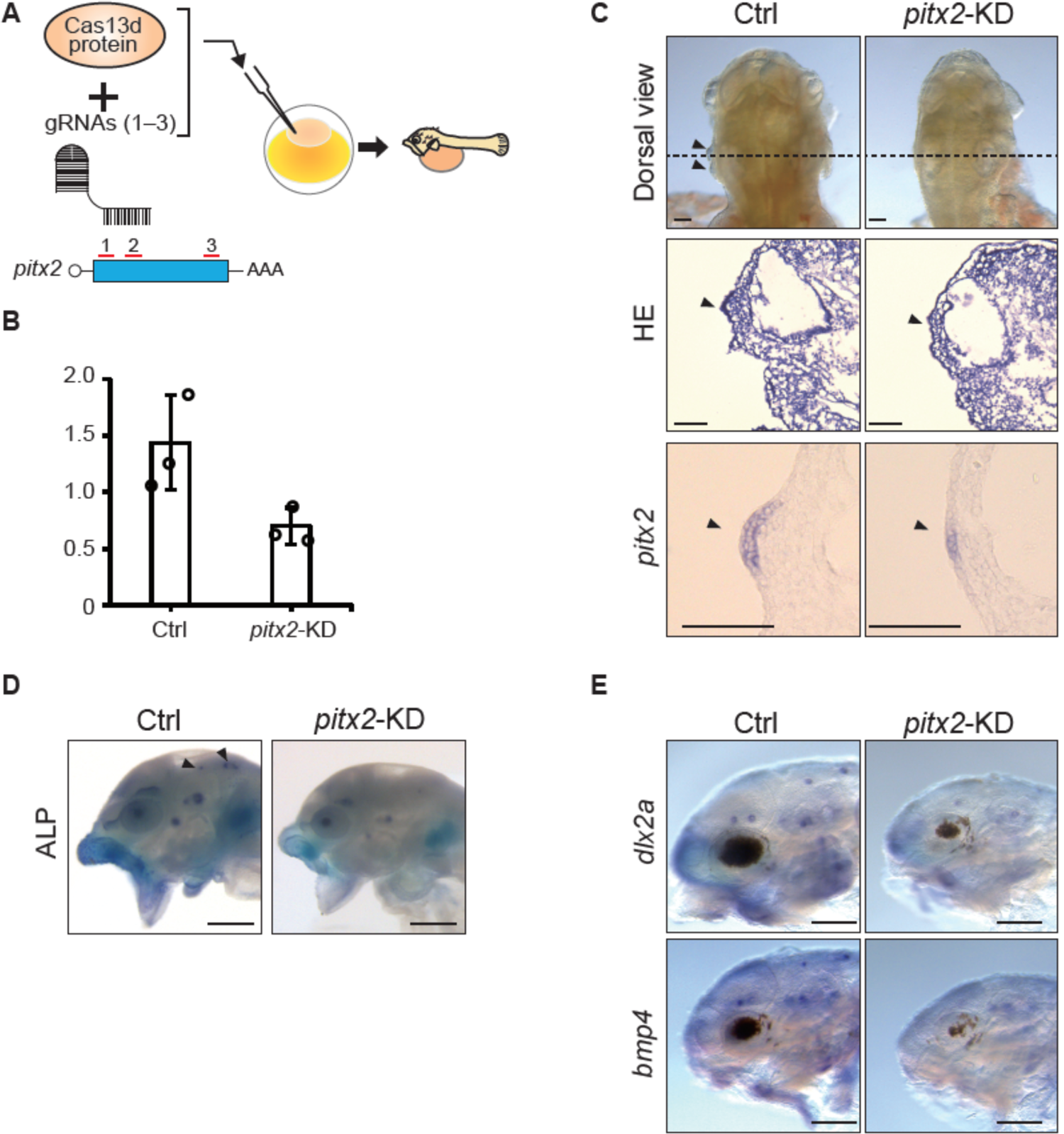
Pitx2 is an inducer of cranial dermal denticle placodes. (A) Schematic diagram of *pitx2* knockdown method for *Ancistrus sp*. embryos using CRISPR-Cas13d. Three gRNA targeting sites in *pitx2* coding sequence are shown by red lines. Cas13d protein was mixed with/without the gRNAs (*pitx2*-KD or Ctrl, respectively) and injected into one- to two-cell stage *Ancistrus sp*. embryos. (B) Quantitative RT-PCR analysis of *pitx2* expression level in Ctrl and *pitx2*-KD embryos at 72 hpf (mean ±s.d, *n*=3, \**P* < 0.05; two-tailed Student’s *t-*test). (C) Representative gross photos of Ctrl (top, left) and *pitx2*-KD (top, right) embryos at 60 hpf. HE staining (middle panels) and *in situ* hybridization of *pitx2* (bottom panels) were conducted using transverse sections at the positions of the dotted lines in top panels. Arrowheads indicate the epithelial placodes. Note that the placode did not thicken in *pitx2*-KD embryos, and *pitx2* expression was downregulated. Scale bars: 100 µm. (D) ALP staining of Ctrl (left) and *pitx2*-KD (right) embryos at 96 hpf. Scale bars: 0.5 mm. (E) Whole-mount *in situ* hybridization of *dlx2a* (top) and *bmp4* (bottom) in Ctrl and *pitx2*-KD embryos at 96 hpf. Scale bars: 0.5 mm. At least three biological replicates were investigated in each experiment.

**Fig. 6.**
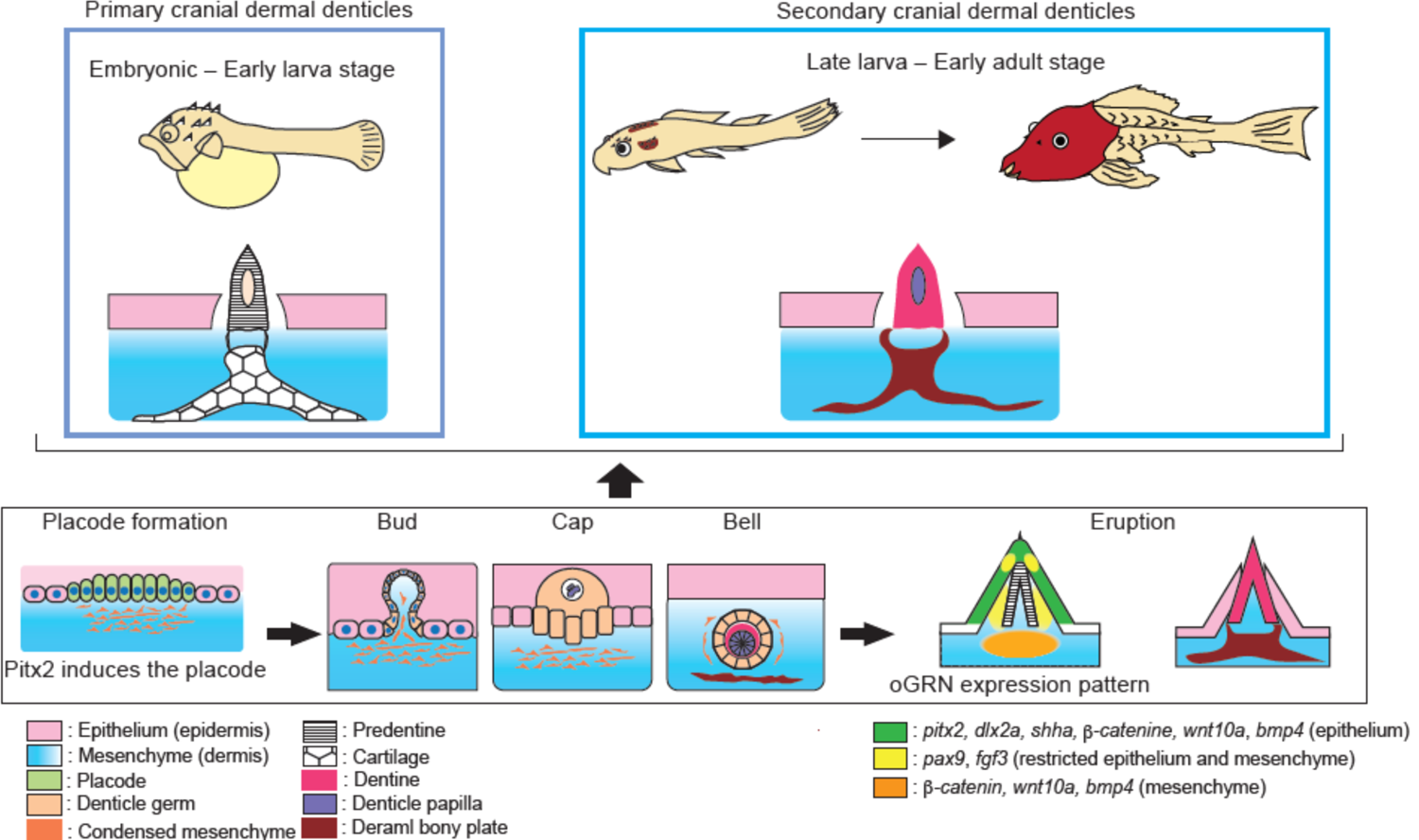
The developmental process of cranial dermal denticles in *Ancistrus sp*. (Top) From the embryonic to the early larva stage (14 dpf), primary cranial dermal denticles form in the restricted positions of the cranial epidermis. After primary cranial dermal denticles shed and disappear, secondary cranial dermal denticles begin to form. The number of secondary cranial dermal denticles increases along with the expansion of underlying dermal bony plates, constructing the head armor. (Bottom) Cranial dermal denticle morphogenesis progresses through the evolutionarily conserved odontode developmental stages. First, the cranial epithelium thickens to form the epithelial placode. The placode invaginates and forms a bud-shaped epithelial layer. Subsequently, condensed mesenchymal cells invade the lumen of the denticle germ, leading to the formation of denticle papilla (bud stage). The denticle germ forms circular epithelial tissue enclosing the denticle papilla (cap stage). The denticle papilla then differentiates into dentin inside the denticle germ, forming a bell-shaped structure (bell stage). At this stage, underlying dermal bony plates differentiate underneath the dental germ. Dentin finally erupts from the epithelium, being anchored to dermal bony plates (eruption stage). The development of primary and secondary cranial dermal denticles is regulated by evolutionarily conserved oGRN. Among oGRN, *pitx2* exhibits the earliest expression in the placode and induces the development of cranial dermal denticles. *Pitx2, dlx2a,* and *shha* are expressed in the epithelial layer surrounding the dentine cone. *Wnt10a, b-catenin,* and *bmp4* are broadly expressed in both the epithelium and underlying mesenchyme. *Pax9* and *fgf3* transcripts were detected in the epithelium and mesenchyme laterally to the dentin cone.

## DISCUSSION

### An evolutionarily conserved dermal odontode developmental process

We comprehensively examined the developmental process of cranial dermal denticles in suckermouth armored catfish. In this system, cranial dermal denticles develop through the histological stages of placode formation, bud, cap, bell, and eruption as previously defined in tooth and placoid scale formation (Balic and Thesleff, 2015; Debiais-Thibaud et al., 2015; Ellis et al., 2015). In particular, the processes of the epithelial folding, dentine differentiation, and dentine eruption closely resemble events previously shown to occur during tooth development. These findings accord with those of a concurrent investigation of a different species of suckermouth armored catfish *Ancistrus triradiatus* (Loricariidae; Siluriformes) (Rivera-Rivera et al., 2021 preprint), and the histological similarities of dermal denticle and tooth formation in both these species implies that dermal denticles in suckermouth armored catfish evolved via redeployment of a preexisting developmental program for teeth in the common ancestor of the catfish family.

### The developmental relationship of dermal denticles and underlying dermal bony plates

Typically, dermal odontodes form anchored to underling dermal bony plates, excepting the placoid scales of cartilaginous fish which form without underlying bony supports (Miyake et al., 1999). Historical hypotheses have proposed that, during dermal odontode development, an underlying dermal bony plate fused with adjacent plates and evolved into large osseous plates. This could be the evolutionary origin of exoskeletons, such as scales or scutes (Reif, 1982; Stensiö, 1962). However, these hypotheses have never been functionally tested due to the lack of developmentally amenable model organisms. In *Ancistrus sp*., we found that primary cranial dermal denticles form anchored to underlying cartilages and secondary cranial dermal denticles develop with underlying dermal bony plates (Fig. 1, and Figs. S1, S2). Intriguingly, the developmental topology of mammalian teeth and the alveolar bone called the tooth-socket is analogous to that of dermal denticles and their underlying support. During tooth development, the dental lamina, which expresses *sema3f* and *fgf8*, induces a mesenchymal condensation that gives rise to the alveolar bone (Diep et al., 2009; Mammoto et al., 2011). Dermal denticles could induce dermal bony plate formation in catfish in an analogous manner considering the similarity of gene expression profiles in tooth and denticle germs (Fig. 2C) (Matalová et al., 2015). Further research is required to understand the developmental relationship between dermal denticles and their underlying dermal bony plates, but dermal denticles could very well be a main driver in regulating dermal bony plate development and divergence of the head armor. However, some Siluriformes catfishes possess dermal bony plates without dermal denticles (Rivera-Rivera and Montoya-Burgos, 2017); thus, the developmental relationship of dermal denticles and basal bony plates could be a derived characteristic gained in the course of Loricariidae fish evolution.

### A conserved oGRN underlies suckermouth armored catfish dermal denticle formation

Our comparative *de novo* RNA-seq analysis identified a conserved oGRN for odontode development in suckermouth armored catfish (Fig. 2), and we found that the expression patterns of representative oGRN genes are strictly conserved in dermal denticles and other odontodes, e.g., teeth and placoid scales (Fig. 3) (Berio and Debiais-Thibaud, 2021; Debiais-Thibaud et al., 2015; Fraser et al., 2010). However, there were some discrepancies. For example, during tooth formation, *pax9* expression is confined to mesenchymal cells, but it is expressed in both the epithelium and mesenchyme of cranial dermal denticles in *Ancistrus sp*. (Berio and Debiais-Thibaud, 2021). Therefore, the epithelial expression of *pax9* could be a derived characteristic in suckermouth armored catfish denticle development.

Notably, the genetic programs of the epithelial placode could have diversified from their evolutionarily ancestral norm. For instance, during mouse taste bud development, which likely retains the ancestral developmental program of the epithelial placode (Fraser et al., 2010), *pax9* is expressed in both the epithelium and mesenchyme (Kist et al., 2014). Teeth might have evolved by a co-option of sensory structure developmental programs (e.g. taste buds) (Bloomquist et al., 2015; Fraser et al., 2010); thus, *pax9* expression might have been lost from the epithelium during tooth evolution. Accordingly, epithelial *pax9* expression could be regained during convergent evolution of dermal denticles in suckermouth armored catfish. Thus, the oGRN might have been prone to recurrent modifications during odontode evolution, producing morphological diversification.

Overall, our *de novo* RNA-seq and ISH results highlight the evolutionarily shared and derived genetic characteristics of dermal odontode development in suckermouth armored catfish. These results suggest that heterotopy and successive modifications of epidermal appendage developmental programs have produced extraordinarily diverse exoskeletal architectures in extinct and living taxa.

### The maturation and mineralization of dermal odontodes in surkermouth armored catfish

Our *de novo* RNA-seq also identified genes likely necessary for maturation and mineralization of dermal denticles. The expression levels of *sparcl1* and *spp1* are higher in juvenile than in adult dermal denticles. *Sparcl1* and *spp1* belong to the secretary calcium binding phosphoprotein (SCPP) family, and *spp1* is thought to have originated from the common ancestral gene *sparcl1* by tandem gene duplication (Kawasaki and Weiss, 2008). *Sparcl1, spp1,* and other genes in the SCPP family exhibit diverse expression patterns in dentine odontoblasts and bone osteoblasts and promote mineralization in various vertebrates. For instance, in vertebrates, *SPARCL1* and the sparcl1-orthologue *SPARC* (also called osteonectin) are exclusively expressed in odontoblasts (Enault et al., 2018; Kawasaki and Weiss, 2008; Liu et al., 2016). However, while fish *spp1* is highly expressed in both odontoblasts and osteoblasts, mammalian *SPP1* is more highly expressed in immature osteoblasts than in odontoblasts (Enault et al., 2018; Kawasaki and Weiss, 2008; Liu et al., 2016). It remains unclear how *sparcl1* and *spp1* function during odontogenesis but they likely contribute to the early mineralization of cranial dermal denticles in suckermouth armored catfish.

In contrast to *sparcl1* and *spp1, β-catenin* and *runx2* are more highly expressed in adult odontodes samples (adult dermal denticles, teeth, and scutes) than in juvenile samples (Figs. 2 C, D and Fig. S3E). The Wnt/β-catenin signaling pathway plays a central role in mouse tooth morphogenesis. In that system, Wnt signaling regulates odontoblast development by stimulating transcription of *runx2*, a gene essential for inducing odontoblast differentiation (Cai et al., 2011; Gaur et al., 2005; Tucker et al., 2000), via direct binding of β-catenin to the *runx2* promoter together with the Tcf1 transcription factor and induces its expression (Gaur et al., 2005). Runx2, in turn, binds to the promoter region of *dspp*, another SCPP family member and indispensable for mineralization, and upregulates its transcription (Chen et al., 2005). Our RNA-seq data indicates that *dspp* transcripts are enriched in adult cranial dermal denticles (Fig. 2D and Fig. S3E) along with *β-catenin* and *runx*2. Thus, a cascade identical or similar to the Wnt/β-catenin–Runx2–DSPP pathway identified in mice likely promotes odontoblast maturation in cranial dermal denticles of suckermouth armored catfish. Our comparative transcriptomic analysis is the first systematic and comprehensive identification of genes involved in the early and late maturation process of dermal odontodes.

### Expression of sox2 in *Ancistrus sp*. primary dermal denticles and teeth indicates regenerative potential

Our *de novo* RNA-seq indicates that dermal odontodes and teeth not only share a significantly conserved oGRN but also share expression of the *SRY-box transcription factor 2* (*sox2*), an epithelial stem cell marker necessary for maintaining progenitors of mammalian teeth epithelial cell lineages (Juuri et al., 2012). We found that *sox2* was abundant in juvenile secondary cranial dermal denticles and adult oral samples but low in adult cranial denticles and scutes (Fig. 2C). Consistent with this, our histological analysis found that *Ancistrus sp*. teeth are polyphyodontic, meaning multiple tooth germs underlie erupted teeth, but successive denticle germs do no underlie adult cranial dermal denticles (Figs. S1C and S2C). Intriguingly, a recent study detected *sox2* transcripts in the dental lamina of shark polyphyodontic teeth, suggesting s*ox2* has an essential function in shark tooth regeneration (Fraser et al., 2019). In contrast, *sox2* transcripts were detected only at the initial developmental stage of placoid scales, suggesting that shark dermal denticles cannot regenerate (Martin et al., 2016). These *sox2* expression patterns in shark teeth and dermal denticles are similar to those of *Ancistrus sp*. Accordingly, we postulate that *Ancistrus sp*. teeth possess the significant regenerative ability, but secondary cranial dermal denticles lose it as *sox2* transcripts decrease during development. It would be intriguing to explore how dermal denticles in suckermouth armored catfish lost *sox2* expression and regenerative ability if they evolved by deploying tooth developmental programs; the odontode developmental programs and regenerative programs could be separable during odontode evolution.

### Redeployment of the tooth developmental program for dermal odontodes development

*Pitx2* expressed in the epithelial placode initiates tooth germ formation in mammals, but *pitx2* expression is undetectable in developing dermal denticles of cartilaginous fish, highlighting a genetic gap in the developmental programs of teeth and dermal odontodes (Debiais-Thibaud et al., 2015). Consistent with the concurrent investigation of the oGRN in another species of suckermouth armored catfish (Rivera-Rivera et al., 2021 preprint), our comparative *de novo* RNA-seq and histological analysis of *Ancistrus sp*. found that *pitx2* was the earliest oGRN gene to be expressed in the epithelial placode of cranial and trunk dermal denticles (Figs. 3, 4 and Fig. S6D). Moreover, the functional knockdown of *pitx2* in suckermouth armored catfish resulted in malformed placodes with decreased expression of *dlx2a* and *bmp4* (Figs. 5 and Fig. S8). These results are consistent with prior studies which showed that reciprocal interactions between Pitx2, Dlx2, and Bmp4 are required to initiate mammalian tooth germ formation (Green et al., 2001; St Amand et al., 2000). Accordingly, we propose that Pitx2 initiates induction of the epithelial placode by triggering oGRN gene expression, including *dlx2* and *bmp4*, in suckermouth armored catfish.

A recent phylogenetic study suggested that Loricarioidae, a catfish suborder including Loricariidae (suckermouth armored catfish) and Callichthyidae (*Corydoras*), independently and repeatedly gained dermal denticles on head and trunk surfaces (Rivera-Rivera and Montoya-Burgos, 2017). However, the underlying genetic mechanisms of this convergent evolution remain elusive. Based on the early expression of *pitx2* and its ability to induce placodes in dermal denticles, we hypothesize that *pitx2* expression might have been redeployed from teeth to dermal denticles in suckermouth armored catfish. Coincidently, the cis-regulatory elements of *pitx1*, another member of *paired-like homeodomain* family, involved in the convergent evolution of fish pelvic fins are under active investigation (Chan et al., 2010; Thompson et al., 2018). It was found that recurrent mutations within the fragile *Pel* element, a specific enhancer of *pitx1,* underlie convergent loss of pelvic fins in sticklebacks (Xie et al., 2019). It is possible that specific regulatory elements of *pitx2* are more susceptible to mutation than regulatory elements of other oGRN genes and this susceptibility contributed to the redeployment of the tooth developmental program into dermal denticles in catfish. Further analysis of dermal denticle evolution in suckermouth armored catfish will help reveal the genetic basis of convergent evolution of dermal odontodes and, possibly, provide us clues to decipher a broader enigma—the evolutionary origins of teeth and dermal odontodes in vertebrate ancestry.

## MATERIALS AND METHODS

### Fish husbandry

Channel catfish (*Ictalurus punctatus*) were commercially purchased from a local fish farm (Live Aquaponics, Florida, USA). Suckermouth armored catfish (*Ancistrus sp*.) were commercially purchased from a local aquarium store and were bred in aquarium tanks. Water was replaced twice a week with dechlorinated water prepared from tap water via chlorine remover treatment (API, tap water conditioner). The water was maintained at 25°C and pH 6.5. After 14 dpf, juveniles and adult fish were fed a boiled vegetables and algae wafer (Tetra) diet. Each tank contained one male and up to two female fish allowing natural breeding. To promote mating, fish caves, which help male fish maintain territories, were installed into the tanks. Males showed typical mating behavior; they repeatedly invited female fish to enter the caves for 3 to 6 hours. After spawning, eggs were transferred into quarantined mesh cages installed in the main tanks, and fries were reared there until 60 dpf. For the following experiments, embryos, juveniles, and adult fish were euthanized with 0.016% MS-222 and fixed as indicated. All animal experiments were approved by the animal committees of Rutgers University (protocol no. 201702646).

### *De novo* RNA sequencing

Mouth, cranial skin, and scute samples were manually dissected from *Ancistrus sp*. juvenile and adult specimens, frozen in liquid nitrogen, and ground into a fine powder by a mortar and pestle. Total RNA was isolated from each powdered sample using the RNase Plus Universal Kit (Qiagen) according to the manufacturer’s instruction. RNA quality assessment, library construction, high-throughput sequencing, and *de novo* transcriptome assembly were conducted by Novogene Co., Ltd (Tianjin, China). Briefly, RNA purity and integrity were measured by an Agilent 2100 Bioanalyzer. Sequencing libraries were constructed with a TruSeq stranded mRNA sample preparation Kit (Illumina), and high-throughput sequencing was carried out to generate paired-end 150 bp reads using the NovaSeq 6000 Reagent Kit (illumina). A total of 1,254,277,785 raw reads were obtained from twelve libraries derived from three replicates each of juvenile cranial dermal denticle (CDj), adult cranial dermal denticle (CD), adult scutes (Scute), and adult teeth (Mouth) samples. The raw reads were filtered to remove adapters, reads containing more than 10% equivocal bases, and reads with base quality less than 20 (Quality score ≤ 20). The clean reads were then *de novo* assembled by Trinity software and 717,406,047 unigenes were identified. Of these unigenes, 488,726 were functionally annotated by aligning them to the Nr (NCBI non-redundant protein sequences, E-value < 1e-5), Nt (NCBI nucleotide sequences, E-value < 1e-5), Pfam (E-value < 0.01), KOG/COG (Eukaryotic Orthologous Groups/Cluster of Orthologous Groups of proteins, E-value < 1e-5), Swiss-Prot (E-value < 1e-5), KEGG (Kyoto Encyclopedia of Genes and Genome, E-value < 1e-5), and GO (Gene Ontology, E-value < 1e-6) databases. Gene functional classification was performed with BLAST2GO based on the GO annotations. Coding sequences (CDS) for unigenes were predicted with BLAST and ESTScasn (3.0.3). To calculate the expression level, the clean reads were mapped against the *de novo* assembled transcriptome using Bowtie. The mapping results of Bowtie were analyzed by RSEM, and the read count and fragments per kilobase of transcript sequence per million mapped reads (FPKM) value were obtained for each sample. The count data was then used as the input for Bioconductor package (3.12) edgeR and DEseq2 to test for differential gene expression. GO enrichment analysis of differentially expressed genes (DEGs) was performed using topGO. Raw reads data is available in the NCBI Sequence Read Archive (SRA) database: PRJNA731927.

### Histological analysis

Bones and cartilage in *Ancistrus sp*. specimens were stained by Alcian blue 8GX (Sigma) and Alizarin red S (Sigma) as previously described (Fernández et al., 2008). Alkaline phosphatase (ALP) staining was performed with BM-purple (Roche) as previously described (Habeck et al., 2002). Paraffin sectioning was performed by the Research Pathology Services of Rutgers University. Briefly, juveniles and adult fish were fixed by 4% PFA at 4° C for one to two weeks and then decalcified in 10% EDTA at 4° C for one to two weeks. After decalcification, samples were dehydrated through a graded ethanol series, cleared in xylene, and embedded in paraffin. Serial sections (8–10 µm) were collected and stained by standard hematoxylin and eosin (HE) staining. Bone-stained samples and HE-stained sections were imaged on an M205 FCA (Leica) stereoscope and Eclipse E800 microscope (Nikon), respectively, equipped with cameras.

### Immunohistochemistry

Fixed samples were washed in PBS, immersed in 15% followed by 30% sucrose in PBS at 4° C, and frozen in OCT embedding compound (Sakura). Frozen sections (8–10 µm) were prepared and treated with an antigen retrieval solution (sodium citrate buffer;10 mM sodium citrate, pH6.0) as previously described (Gu et al., 2016). Sections were then permeabilized in 0.3% Triton X-100 in PBS for 15 min and blocked in blocking buffer (5% sheep serum and 0.1% Tween-20 in PBS) for 1 hour. The PCNA (CST) antibody was diluted in blocking buffer (1:500) and incubated with the sections overnight. Then, sections were washed in 0.1% Tween-20 in PBS and incubated with a secondary antibody (Alexa Fluor 488, Molecular Probes) in blocking buffer (1:1000) for 1 hour at room temperature (RT). Counter nuclear staining was performed with DAPI in blocking buffer (1:2000). Images were captured with the LSM 510 Meta inverted confocal microscope (Zeiss).

### *in situ* hybridization

Based on the CDS prediction for the *de novo* transcriptome, cloning primers (Table. S1) were designed to amplify RNA probes from cDNA. Probes were inserted into the pCR-Blunt II-TOPO vector or pCRII-TOPO vector (Invitrogen). DIG-labeled RNA probes were synthesized with the Riboprobe Systems (Promega) and DIG RNA Labeling Mix (Roche). Whole mount *in situ* hybridization (ISH) was performed as described (O’Neill et al., 2007). Before rehydration, 30 dpf juveniles were treated with 6% hydrogen peroxide in methanol for 30 min at RT. After rehydration, embryos and juveniles were treated with Proteinase-K (10 µg/ml) in PBS for 15 min at RT and 30 min at 37° C, respectively. Samples were fixed with 0.5% glutaraldehyde plus 4% PFA in PBS for 20 min at RT and then the hybridization was performed.

Paraffin and frozen sections were prepared as described above and subjected to ISH as previously described (Braissant and Wahli, 1998). Paraffin sections were rehydrated (or frozen sections washed in PBS) followed by permeabilization in 0.1% Triton X-100 in PBS for 30 min at RT. Tissue sections were equilibrated in 0.1 M triethanolamine with 0.25% acetic anhydride and were hybridized with an RNA probe at 70° C overnight. The signal was detected with an anti-DIG-AP Fab fragment (Roche) and BM-purple, and images of whole mount ISH or section ISH were captured with an M205 FCA (Leica) stereoscope or Eclipse E800 microscope (Nikon), respectively.

### Designing of guide RNA targeting pitx2 and preparation of gRNA-Cas13d solution

CRISPR-RfxCas13d knockdown was performed as described previously (Kushawah et al., 2020). Briefly, the pET-28b-RfxCas13d-His vector (Addgene, Plasmid #141322) was used for RfxCas13d protein production (Bon Opus Biosciences, LLC). The final concentration of RfxCas13d was adjusted to 3 µg/µl in storage buffer (50 mM HEPES-KOH, pH 7.5, 250 mM KCl, 1 mM DTT, 10% Glycerol) and stored at −80°C. For gRNA synthesis, the *Ancistrus* Pitx2 CDS was analyzed for high accessibility sites using RNAfold software (http://rna.tbi.univie.ac.at//cgi-bin/RNAWebSuite/RNAfold.cgi) and identified 22 nucleotides to generate each guide RNA (gRNA). A gRNA universal forward primer containing a T7 promoter and reverse primers containing target sites are shown in Table. S1. gRNA templates were generated with PCR using a pool of 3 different gRNA primers at equal concentrations and were *in vitro* transcribed using the mMESSAGE T7 Kit (Invitrogen). The final concentration of gRNAs was adjusted to 1600 ng/µl.

### Suckermouth catfish microinjection

To generate the ribonucleoproteins (RNPs), 1µl of gRNA (1600 ng/µl) and 4.5 µl of RfxCas13d protein (3 mg/ml) were mixed in 1 µl of nuclease-free water with phenol red (Invitrogen). One to 5 nL of RNPs were injected into one- to two-cell stage *Ancistrus sp*. embryos according to the zebrafish standard method. Briefly, fertilized eggs were transferred to a petri dish and washed with distilled water. Eggs that stuck together were carefully detached using fine tweezers. After RNP injection, embryos were incubated for 24 h in 0.0001% methylene blue plus distilled water in a 28° C incubator with aeration and then were transferred into a mesh cage installed in the main tanks.

### qRT-PCR

Total RNA was purified from Ctrl and *pitx2*-KD embryos at 72 hpf using the RNeasy Plus Universal Kit (QIAGEN), and cDNA synthesis was performed using the iScript cDNA Synthesis Kit (Bio-Rad). qRT-PCR was performed with a SYBR Green PCR Master Mix and the 7900 Real time PCR system (Applied Biosystems). Results were analyzed by the standard curve method and normalized to β-actin expression. Three biological replicates were investigated in this experiment. Primer sequences are shown in Table. S1.

## Acknowledgements

We would like to thank Dr. Erin Kelly (Rutgers) for critical comments, which substantially improved this paper.

## Author contributions

S.M. performed research. S.M. and T.N. analyzed the data and wrote the paper.

## Funding

This work was supported by the TOYOBO Biotechnology Foundation, The Uehara Memorial Foundation, and Mochida Memorial Foundation for Medical and Pharmaceutical Research Fellowship (to S.M.); institutional support was provided by the Rutgers University School of Arts and Sciences and the Human Genetics Institute of New Jersey (to T.N.).

## Data availability

*De novo* RNA sequencing assembly data have been deposited in the National Center for Biotechnology Information Sequence Read Archives (NCBI SRA), www.ncbi.nlm.nih.gov/sra (BioProject accession code <u>PRJNA288370</u>).

## Competing interests

The authors have declared no competing interest.

## Supplementary Figures/table

**Fig. S1.**
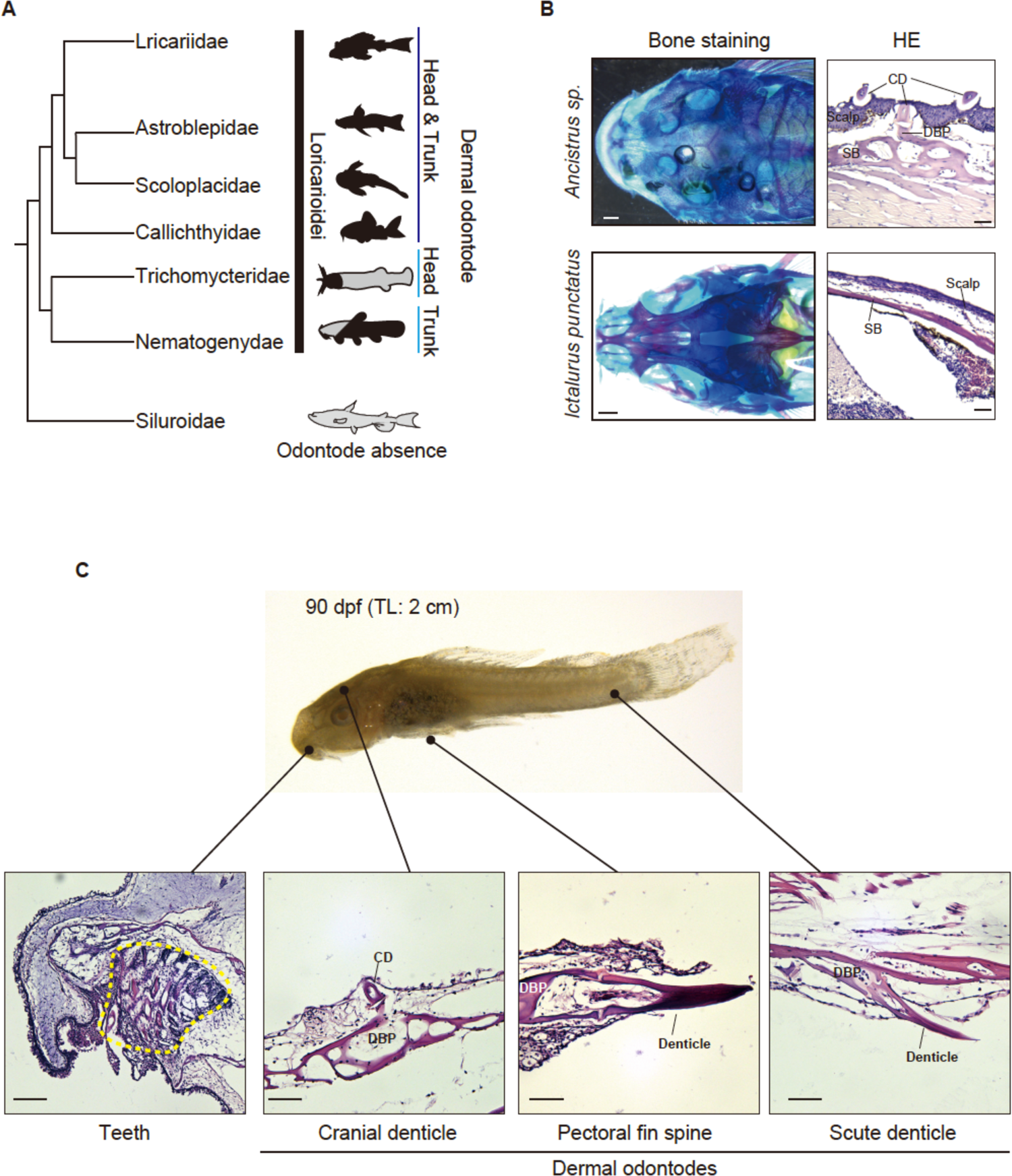
Formation of dermal odontodes in *Ancistrus sp*. A) Phylogenetic tree of Siluriforms. Tree topology from Rivera-Rivera and Montoya-Burgo, 2017. Loricariidae, Astroblepidae, Scoloplacidae, and Calichythyidae form dermal odontodes in the whole-body surface (black). Astroblepidae and Trichomycteridae form dermal odontodes in the head and trunk, respectively. By contrast, Siluroidae lacks dermal odontodes (gray). (B) Bone staining pictures and HE-stained transverse sections of suckermouth armored catfish (*Ancistrus sp.,* TL: 6 cm, top, n=2) and channel catfish (*Ictalurus punctatus*, TL: 5 cm, bottom, n=6). (C) Representative pictures of *Ancsitrus sp.* juvenile at 90 dpf (TL: 2 cm, top) and HE-stained sagittal sections of oral tissue, cranial dermal denticles, pectoral fin spines, and scute denticles (bottom panels). Two biological replicates were investigated in this experiment. The yellow dotted area indicates teeth. Scale bar: 0.5 mm. CD; cranial dermal denticle, SB; skull bone, DBP; dermal bony plate.

**Fig. S2.**
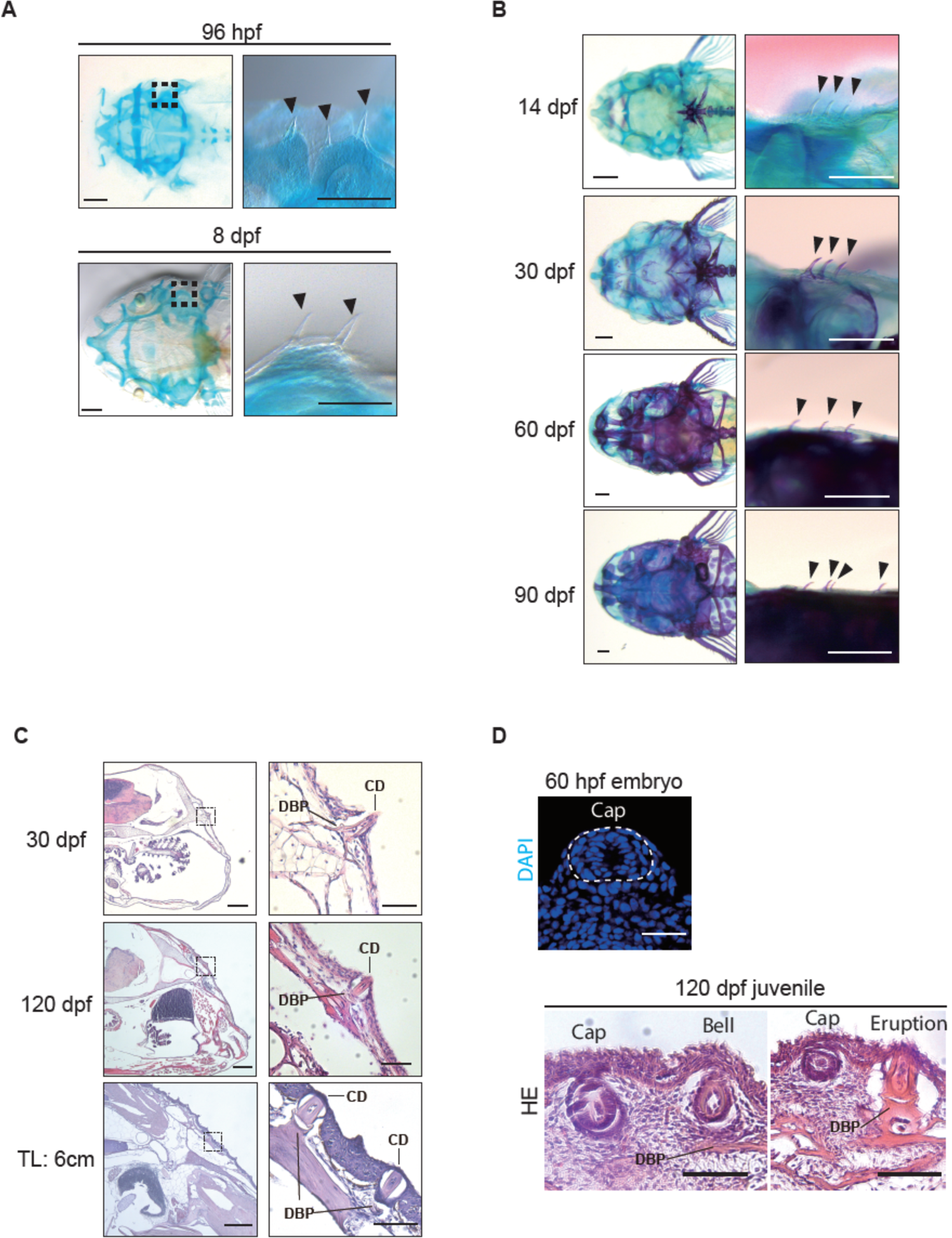
Whole-mount and histological analysis of developing cranial dermal denticles from the embryonic to the adult stage. (A) Bone and cartilage staining of the developmental series of suckermouth armored catfish embryos at 96 hpf (left) and larva at 8 dpf (right). Each right panel shows higher magnification of the dotted box area in the left panels. Arrowheads indicate the primary cranial dermal denticles. Scale bars: 0.5 mm. (B) Bone and cartilage staining of the developmental series of juveniles at 14, 30, 60, and 90 dpf. Each right picture shows high-magnification view of the secondary cranial dermal denticle formation sites (arrowheads) in the left pictures. Scale bar: 0.5 mm. (C) HE-stained head transverse sections of the juveniles at 30 dpf, 120 dpf, and the early adult stage (TL: 6 cm). Each right panel shows high-magnification view of the dotted box area in the left panels. Scale bars: 0.5 mm (left) and 0.1 mm (right). (D) Nuclear staining (DAPI) of a transverse section of a primary cranial dermal denticle at the cap stage in 60 hpf embryo (top panel). HE-staining of transverse sections of 120 dpf juvenile (TL: 3 cm, bottom panels). Secondary cranial dermal denticles form in the cranial epidermis through the cap, bell, and eruption stages. Scale bars: 100 µm. CD; cranial dermal denticle, DBP; dermal bony plate. At least two biological replicates were investigated in each experiment.

**Fig. S3.**
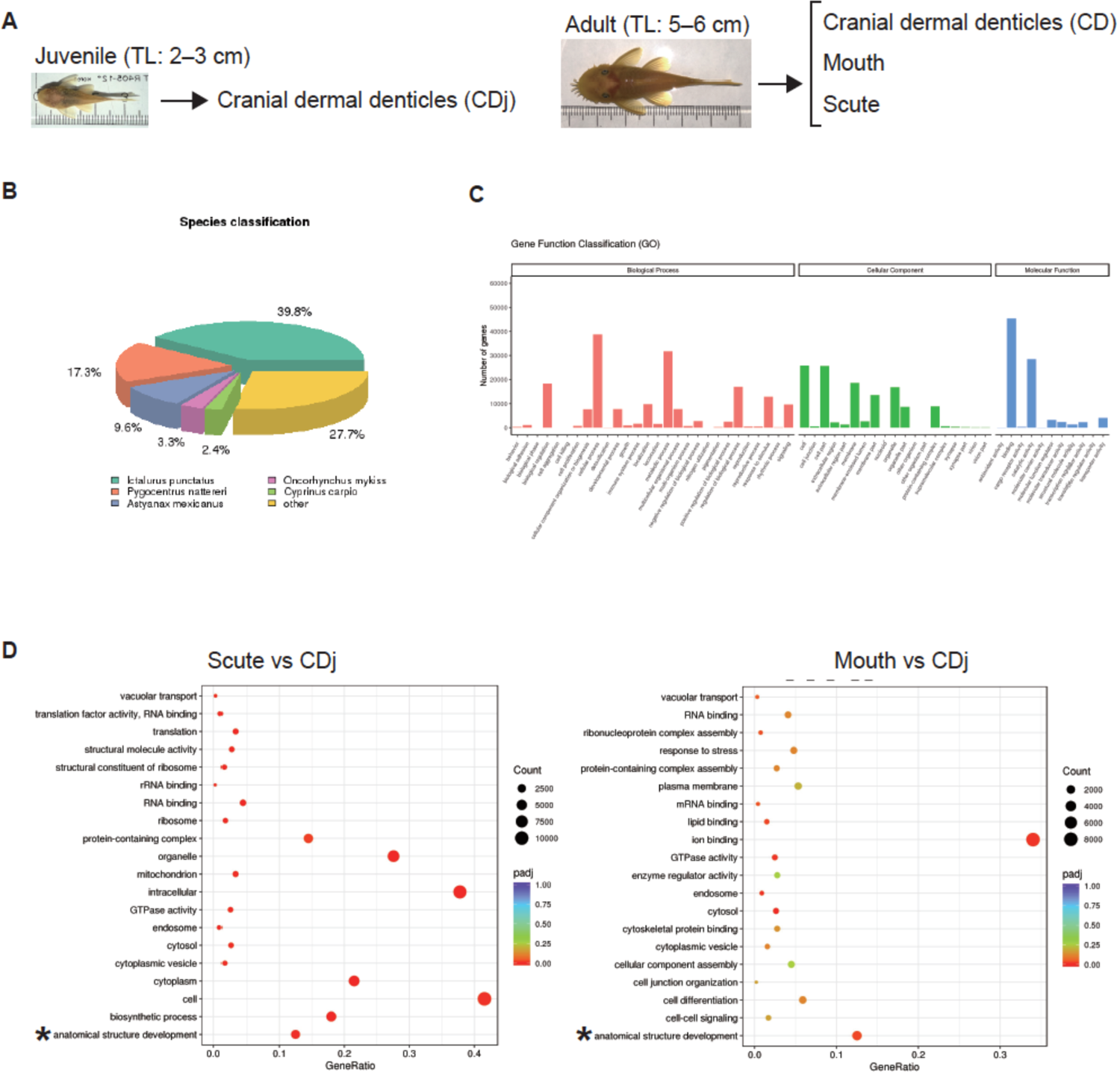
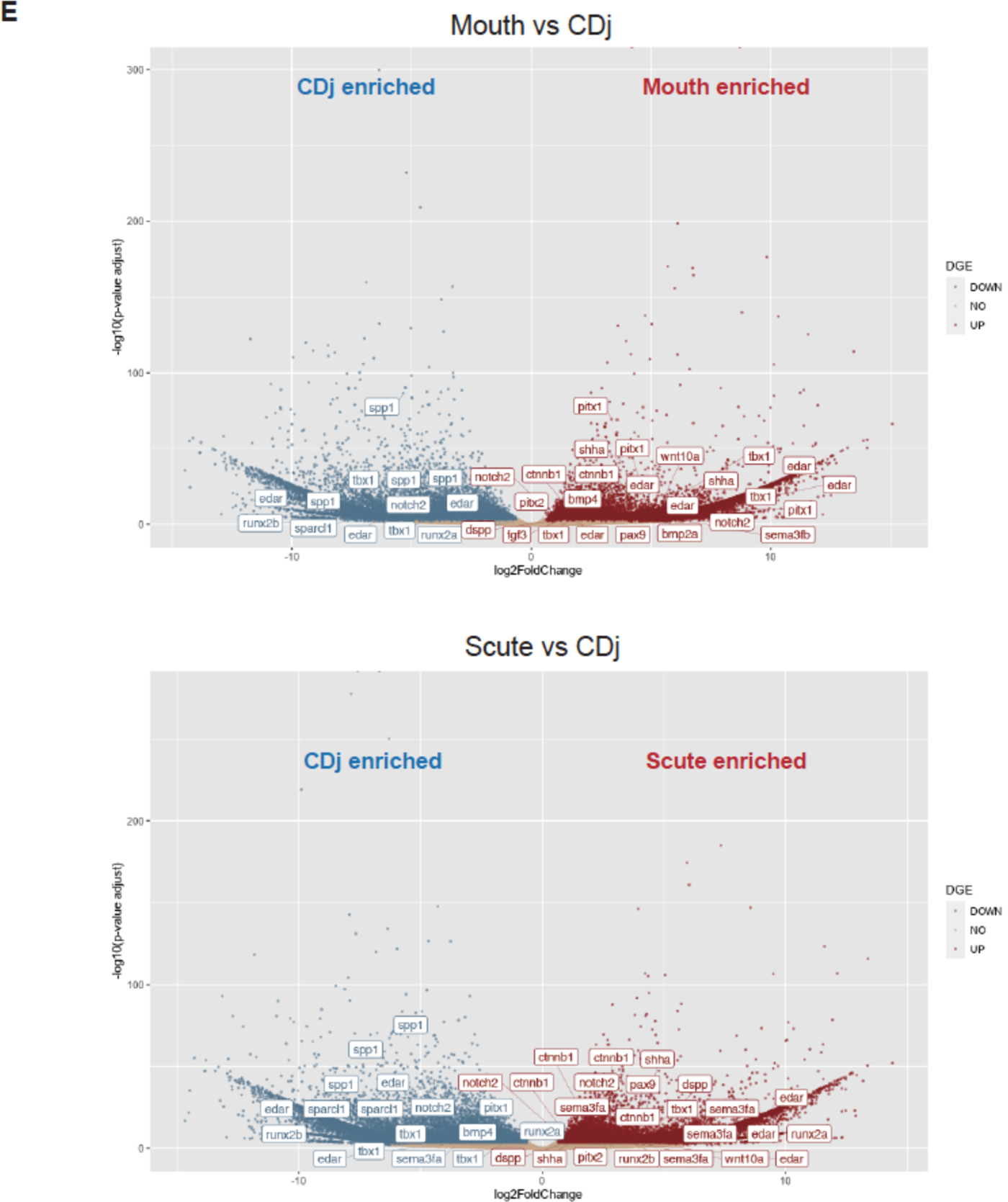
*De novo* transcriptome profiling in the odontode tissues of *Ancistrus sp*. (A) *De novo* RNA-seq was performed by using odontode tissues collected from juvenile cranial skins (left, TL: 2–3 cm, *n*=3) and adult cranial skins, mouths, and scutes (right, TL: 5– 6 cm, *n*=3). (B) Top-hit species distribution based on Nr-annotated unigenes. (C) Functional gene classification of GO-annotated unigenes. (D) GO enrichment analysis of the enriched genes in mouths (left) and scutes (right), compared to CDj. The vertical axis represents the GO terms and the horizontal axis does the enrichment of each term. The size of each point represents the number of upregulated genes in the GO terms, and the color of the point represents the adjusted *p*-value. (E) The volcano plots display the enriched genes in Mouth (top) and Scute (bottom) relative to CDj. Significantly enriched genes in CD are highlighted in red (log2foldchange > 0.6 and adjusted *p*-value < 0.01) and enriched genes in CDj are in blue (log2foldchange < −0.6 and adjusted *p*-value < 0.01). The gene symbols show differentially expressed genes associated with oGRN and odontogenesis. As different unigenes were annotated by the same gene names or homologous genes in different species, the multiple same gene names were shown in the plot.

**Fig. S4.**
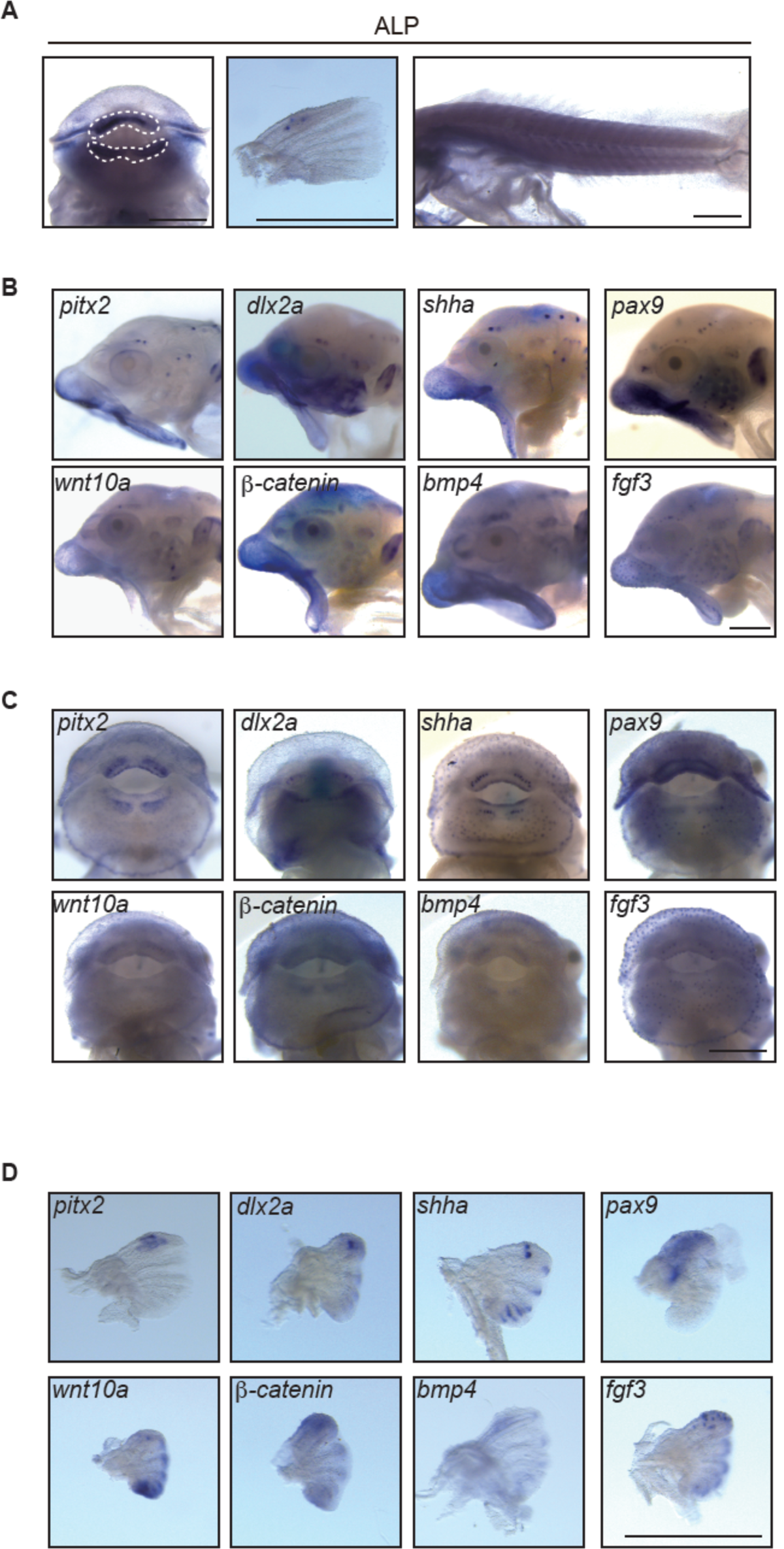
Expression of oGRN genes in the odontode tissues of *Ancsitrus sp.* (A) ALP staining of teeth (left, white dotted outline), pectoral fins (middle), and the trunk (right) of *Ancistrus sp*. embryos at 96 hpf. Teeth and pectoral fin spines, not the trunk, showed restricted ALP staining. (B–D) Whole-mount *in situ* hybridization of *pitx2, dlx2a, shha, pax9, wnt10a, b-catenin, bmp4,* and *fgf3*. All of these gene transcripts were detected in the cranial dermal denticles (B), teeth (C), and pectoral fin spines (D). Scale bars: 0.5 mm. At least three biological replicates were investigated in each experiment.

**Fig. S5.**
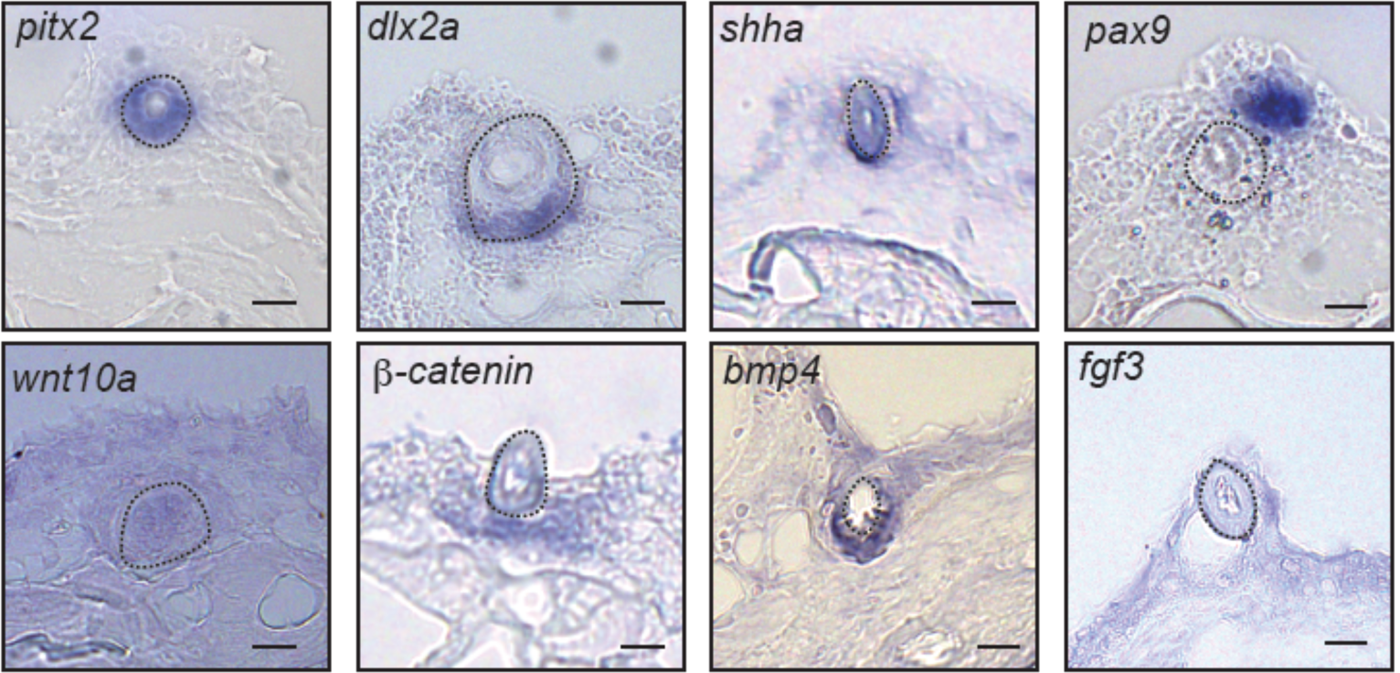
Expression of oGRN genes in the secondary cranial dermal denticles. Transverse sections of whole-mount *in situ* hybridization of *pitx2, dlx2a, shha, pax9, wnt10a, b-catenin, bmp4,* and *fgf3* in juvenile (120 dpf, TL: 3 cm) cranial dermal denticles. Scale bars: 100 µm. Dotted circles indicate denticle germ. Note that, to display the representative oGRN gene expression patterns, we selected different slice positions of dermal denticles for some of the section ISH photos. Accordingly, some denticle germ morphology appear to be different from others. These data were obtained from at least two biological replicates.

**Fig. S6.**
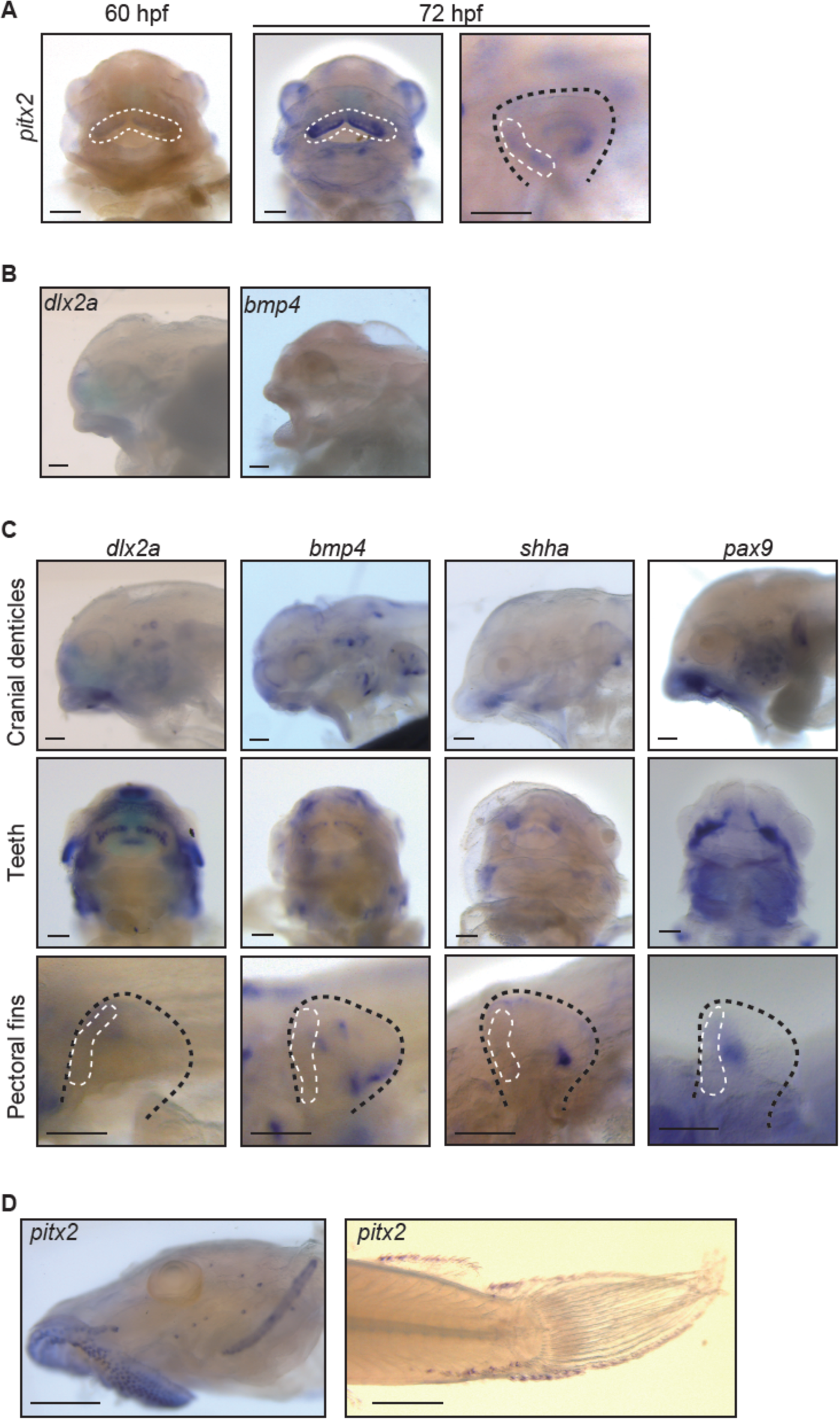
*Pitx2* is expressed at the initiation stage of cranial dermal denticle development. (A) Whole-mount *in situ* hybridization of *pitx2* in *Ancistrus sp*. embryo at 60 hpf (teeth) and 72 hpf (left: teeth and right: pectoral fin, black dotted outline). The expression of *pitx2* was initiated in teeth (white dotted circles) and pectoral fin spines (white dotted circle) from 60 hpf and 72 hpf, respectively. Scale bars: 100 µm. (B) Whole-mount *in situ* hybridization of *dlx2a* and *bmp4* in *Ancistrus sp*. at 60 hpf. *Dlx2a* and *bmp4* transcripts were not detected. Scale bars: 100 µm. (C) Whole-mount *in situ* hybridization of *dlx2a, bmp4, shha,* and *pax9* in cranial dermal denticles (top), teeth (middle), and pectoral fin spines (bottom, black dotted outline) of *Ancistrus sp*. embryos at 72 hpf. Scale bars: 100 µm. *Dlx2a* and *bmp4* but not *shha* and *pax9* transcripts were detected in the cranial dermal denticles and teeth germs at 72 hpf. At this stage, all of these genes are not expressed at the prospective spine denticle sites (white dotted circles). (D) Whole-mount *in situ* hybridization of *pitx2* in *Ancistrus sp*. juvenile at 30 dpf. The expression of *pitx2* was detected at newly forming cranial dermal denticles in the head (left) and the trunk (right). Scale bars: 0.5 mm. At least three biological replicates were investigated in each experiment.

**Fig. S7.**
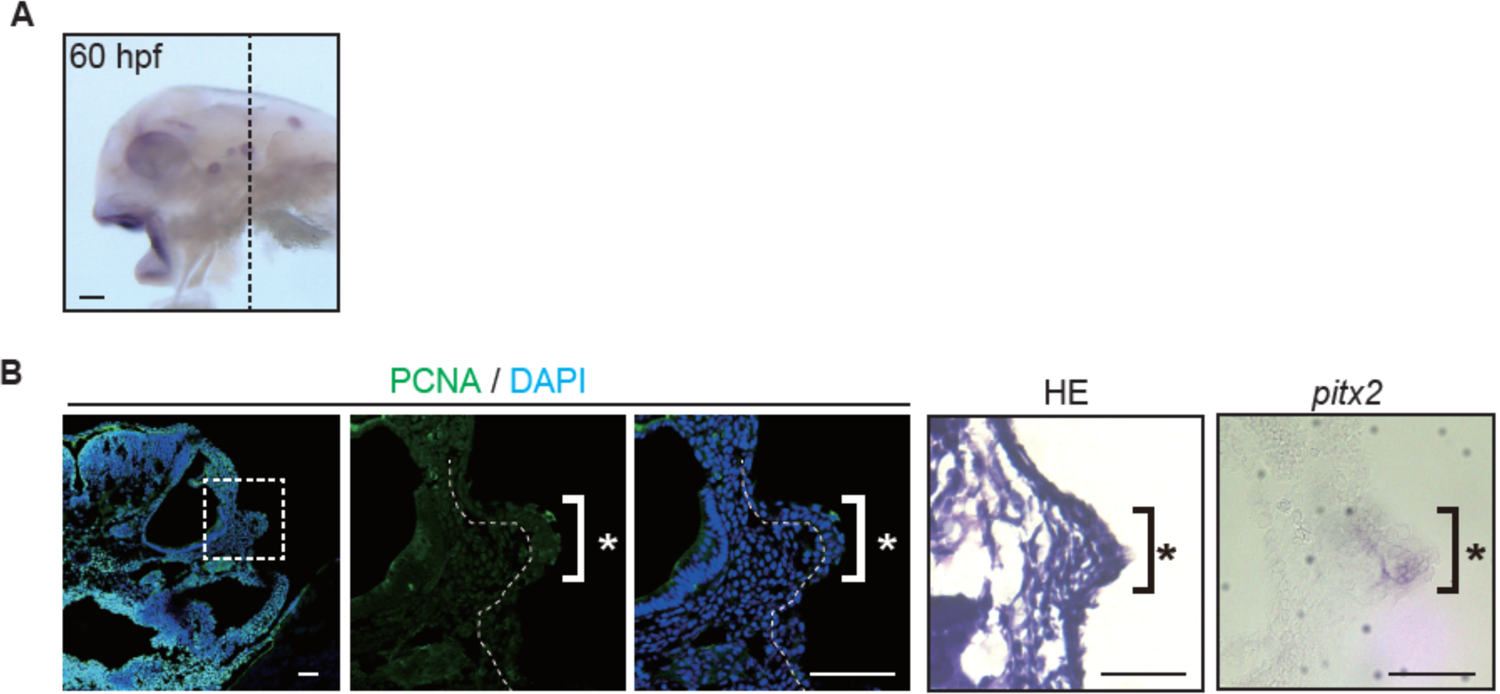
Formation of the cranial dermal denticle placodes in *Ancistrus sp*. embryos. (A) Whole-mount *in situ* hybridization of *pitx2* in an *Ancistrus sp*. embryo at 60 hpf. Scale bars: 100 µm. (B) Immunostaining of PCNA with nuclear staining (DAPI), HE staining, and *in situ* hybridization of *pitx2* for transverse sections at the position of the dotted line in A. *Pitx2* transcripts were detected in the anatomically defined placode (asterisk; thickened epithelial layer with the reduction of cell proliferation). Scale bars: 50 µm. At least three biological replicates were investigated in each experiment.

**Fig. S8.**
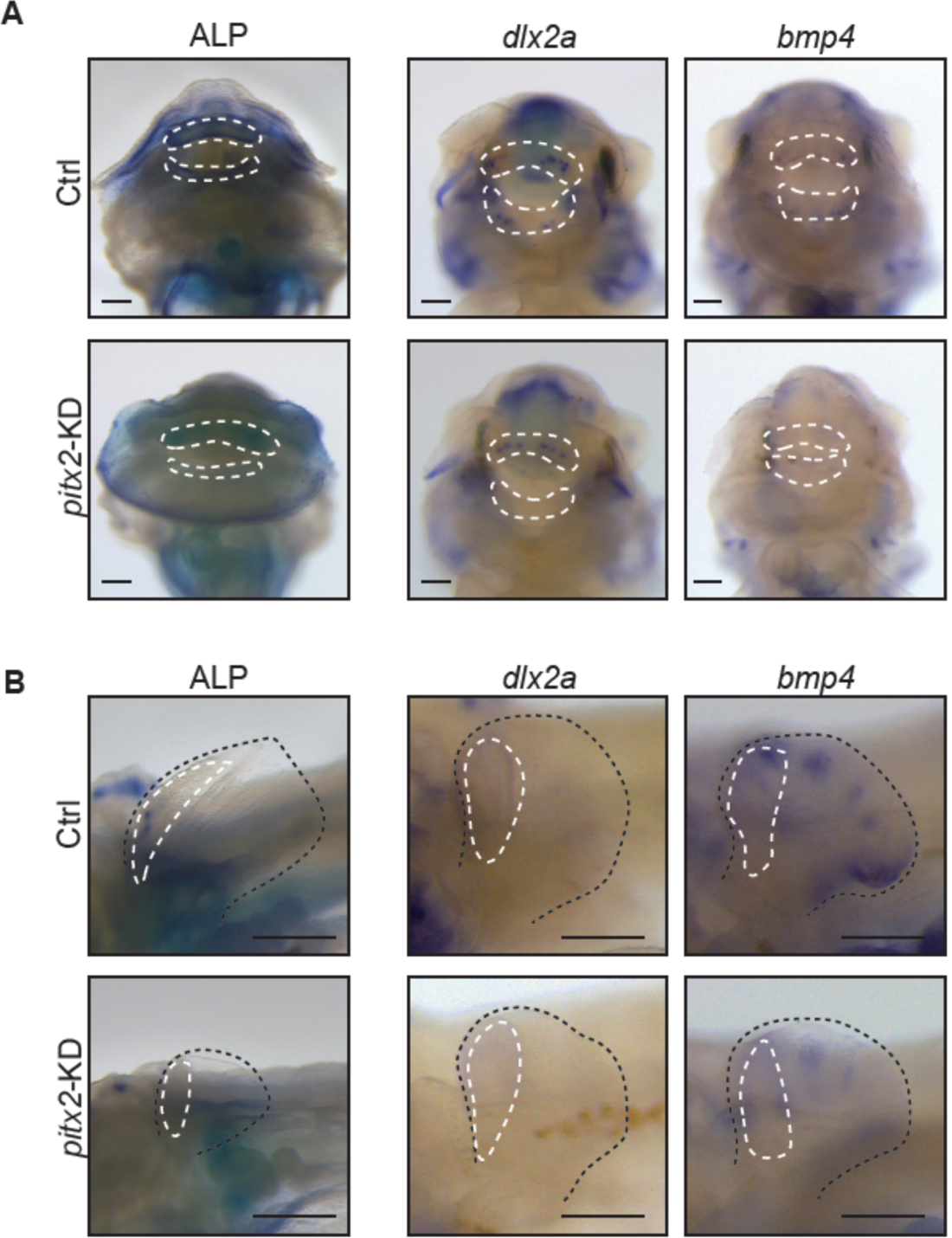
Knockdown of *pitx2* reduces teeth and pectoral fin spines formation. (A, B) Embryos injected with only Cas13d protein (Ctrl)- or Cas13d protein with *pitx2*-gRNAs (*pitx2*-KD) at 96 hpf. Knockdown of *pitx2* reduced the ALP staining and the expression of *dlx2a* and *bmp4* in teeth (A, white dotted circles, scale bars: 0.5 mm) and pectoral fin spines (B, white dotted circles, Scale bars: 100 µm.). Dotted black outlines indicate the pectoral fins in B. At least three biological replicates were investigated in each experiment.

**Table. S1.**
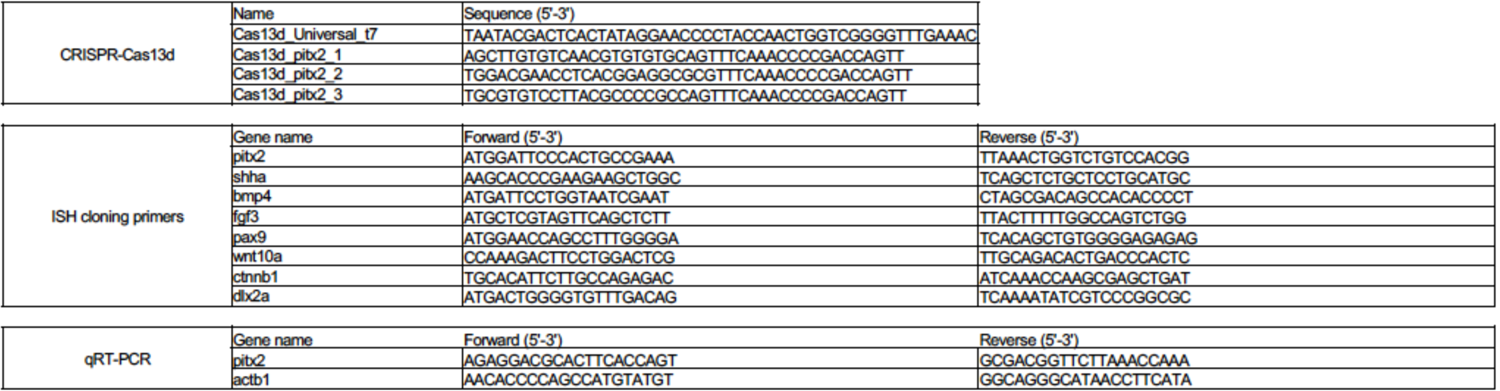
Cas13d oligonucleotides and primers list.

## References

1. Alexander, R. M. (1966). Structure and function in the catfish. Journal of Zoology 148, 88–152.

2. Balic, A. and Thesleff, I. (2015). Tissue Interactions Regulating Tooth Development and Renewal. Curr Top Dev Biol 115, 157–186.

3. Bei, M., Kratochwil, K. and Maas, R. L. (2000). BMP4 rescues a non-cell-autonomous function of Msx1 in tooth development. Development 127, 4711–4718.

4. Berio, F. and Debiais-Thibaud, M. (2021). Evolutionary developmental genetics of teeth and odontodes in jawed vertebrates: a perspective from the study of elasmobranchs. J Fish Biol 98, 906–918.

5. Bitgood, M. J. and McMahon, A. P. (1995). Hedgehog and Bmp genes are coexpressed at many diverse sites of cell-cell interaction in the mouse embryo. Dev Biol 172, 126–138.

6. Bloomquist, R. F., Parnell, N. F., Phillips, K. A., Fowler, T. E., Yu, T. Y., Sharpe, P. T. and Streelman, J. T. (2015). Coevolutionary patterning of teeth and taste buds. Proc Natl Acad Sci U S A 112, E5954–5962.

7. Braissant, O. and Wahli, W. (1998). A Simplified In Situ Hybridization Protocol Using Non-redioactively Labeled Probes to Detect Abundant and Rare mRNAs on Tissue Sections. BIOCHEMICA s *NO* 1, 10–16.

8. Cai, X., Gong, P., Huang, Y. and Lin, Y. (2011). Notch signalling pathway in tooth development and adult dental cells. Cell Prolif 44, 495–507.

9. Chan, Y. F., Marks, M. E., Jones, F. C., Villarreal, G., Jr., Shapiro, M. D., Brady, S. D., Southwick, A. M., Absher, D. M., Grimwood, J., Schmutz, J., et al. (2010). Adaptive evolution of pelvic reduction in sticklebacks by recurrent deletion of a Pitx1 enhancer. Science 327, 302–305.

10. Chen, J., Lan, Y., Baek, J. A., Gao, Y. and Jiang, R. (2009). Wnt/beta-catenin signaling plays an essential role in activation of odontogenic mesenchyme during early tooth development. Dev Biol 334, 174–185.

11. Chen, S., Rani, S., Wu, Y., Unterbrink, A., Gu, T. T., Gluhak-Heinrich, J., Chuang, H. H. and Macdougall, M. (2005). Differential regulation of dentin sialophosphoprotein expression by Runx2 during odontoblast cytodifferentiation. J Biol Chem 280, 29717–29727.

12. Cooper, R. L., Martin, K. J., Rasch, L. J. and Fraser, G. J. (2017). Developing an ancient epithelial appendage: FGF signalling regulates early tail denticle formation in sharks. Evodevo 8, 8.

13. Cooper, R. L., Thiery, A. P., Fletcher, A. G., Delbarre, D. J., Rasch, L. J. and Fraser, G. J. (2018). An ancient Turing-like patterning mechanism regulates skin denticle development in sharks. Sci Adv 4, eaau5484.

14. Dassule, H. R. and McMahon, A. P. (1998). Analysis of epithelial-mesenchymal interactions in the initial morphogenesis of the mammalian tooth. Dev Biol 202, 215–227.

15. Debiais-Thibaud, M., Chiori, R., Enault, S., Oulion, S., Germon, I., Martinand-Mari, C., Casane, D. and Borday-Birraux, V. (2015). Tooth and scale morphogenesis in shark: an alternative process to the mammalian enamel knot system. BMC Evol Biol 15, 292.

16. Debiais-Thibaud, M., Oulion, S., Bourrat, F., Laurenti, P., Casane, D. and Borday-Birraux, V. (2011). The homology of odontodes in gnathostomes: insights from Dlx gene expression in the dogfish, Scyliorhinus canicula. BMC Evol Biol 11, 307.

17. Di-Poï, N. and Milinkovitch, M. C. (2016). The anatomical placode in reptile scale morphogenesis indicates shared ancestry among skin appendages in amniotes. Sci Adv 2, e1600708.

18. Diep, L., Matalova, E., Mitsiadis, T. A. and Tucker, A. S. (2009). Contribution of the tooth bud mesenchyme to alveolar bone. J Exp Zool B Mol Dev Evol 312b, 510-517.

19. Ellis, N. A., Glazer, A. M., Donde, N. N., Cleves, P. A., Agoglia, R. M. and Miller, C. T. (2015). Distinct developmental genetic mechanisms underlie convergently evolved tooth gain in sticklebacks. Development 142, 2442–2451.

20. Enault, S., Muñoz, D., Simion, P., Ventéo, S., Sire, J. Y., Marcellini, S. and Debiais-Thibaud, M. (2018). Evolution of dental tissue mineralization: an analysis of the jawed vertebrate SPARC and SPARC-L families. BMC Evol Biol 18, 127.

21. Fernández, I., Hontoria, F., Ortiz-Delgado, J. B., Kotzamanis, Y., Estévez, A., Zambonino-Infante, J. L. and Gisbert, E. (2008). Larval performance and skeletal deformities in farmed gilthead sea bream (Sparus aurata) fed with graded levels of Vitamin A enriched rotifers (Brachionus plicatilis). Aquaculture 283, 102–115.

22. Fierstine, H. L. (1990). A Paleontological Review of Three Billfish Families (Istiophoridae, Xiphiidae, and Xiphiorhynchidae).

23. Fraser, G. J., Cerny, R., Soukup, V., Bronner-Fraser, M. and Streelman, J. T. (2010). The odontode explosion: the origin of tooth-like structures in vertebrates. Bioessays 32, 808–817.

24. Fraser, G. J., Hamed, S. S., Martin, K. J. and Hunter, K. D. (2019). Shark tooth regeneration reveals common stem cell characters in both human rested lamina and ameloblastoma. Sci Rep 9, 15956.

25. Gaur, T., Lengner, C. J., Hovhannisyan, H., Bhat, R. A., Bodine, P. V., Komm, B. S., Javed, A., van Wijnen, A. J., Stein, J. L., Stein, G. S., et al. (2005). Canonical WNT signaling promotes osteogenesis by directly stimulating Runx2 gene expression. J Biol Chem 280, 33132–33140.

26. Green, P. D., Hjalt, T. A., Kirk, D. E., Sutherland, L. B., Thomas, B. L., Sharpe, P. T., Snead, M. L., Murray, J. C., Russo, A. F. and Amendt, B. A. (2001). Antagonistic regulation of Dlx2 expression by PITX2 and Msx2: implications for tooth development. Gene Expr 9, 265–281.

27. Gu, L., Cong, J., Zhang, J., Tian, Y. Y. and Zhai, X. Y. (2016). A microwave antigen retrieval method using two heating steps for enhanced immunostaining on aldehyde-fixed paraffin-embedded tissue sections. Histochem Cell Biol 145, 675–680.

28. Habeck, H., Odenthal, J., Walderich, B., Maischein, H. and Schulte-Merker, S. (2002). Analysis of a zebrafish VEGF receptor mutant reveals specific disruption of angiogenesis. Curr Biol 12, 1405–1412.

29. Haspel, G., Schwartz, A., Streets, A., Camacho, D. E. and Soares, D. (2012). By the teeth of their skin, cavefish find their way. Curr Biol 22, R629–630.

30. Huysseune, A. and Sire, J. Y. (1998). Evolution of patterns and processes in teeth and tooth-related tissues in non-mammalian vertebrates. Eur J Oral Sci 106 Suppl 1, 437–481.

31. Janvier, P. (1996). *Early vertebrates*: Oxford University Press.

32. Juuri, E., Saito, K., Ahtiainen, L., Seidel, K., Tummers, M., Hochedlinger, K., Klein, O. D., Thesleff, I. and Michon, F. (2012). Sox2+ stem cells contribute to all epithelial lineages of the tooth via Sfrp5+ progenitors. Dev Cell 23, 317–328.

33. Kawasaki, K., Suzuki, T. and Weiss, K. M. (2004). Genetic basis for the evolution of vertebrate mineralized tissue. Proc Natl Acad Sci U S A 101, 11356–11361.

34. Kawasaki, K. and Weiss, K. M. (2008). SCPP gene evolution and the dental mineralization continuum. J Dent Res 87, 520–531.

35. Keating, J. N. and Donoghue, P. C. (2016). Histology and affinity of anaspids, and the early evolution of the vertebrate dermal skeleton. Proc Biol Sci 283, 20152917.

36. Keating, J. N., Marquart, C. L. and Donoghue, P. C. (2015). Histology of the heterostracan dermal skeleton: Insight into the origin of the vertebrate mineralised skeleton. J Morphol 276, 657–680.

37. Kist, R., Watson, M., Crosier, M., Robinson, M., Fuchs, J., Reichelt, J. and Peters, H. (2014). The formation of endoderm-derived taste sensory organs requires a Pax9-dependent expansion of embryonic taste bud progenitor cells. PLoS Genet 10, e1004709.

38. Komori, T. (2010). Regulation of bone development and extracellular matrix protein genes by RUNX2. Cell Tissue Res 339, 189–195.

39. Kushawah, G., Hernandez-Huertas, L., Abugattas-Nuñez Del Prado, J., Martinez-Morales, J. R., DeVore, M. L., Hassan, H., Moreno-Sanchez, I., Tomas-Gallardo, L., Diaz-Moscoso, A., Monges, D. E., et al. (2020). CRISPR-Cas13d Induces Efficient mRNA Knockdown in Animal Embryos. Dev Cell 54, 805–817.e807.

40. Li, S., Kong, H., Yao, N., Yu, Q., Wang, P., Lin, Y., Wang, J., Kuang, R., Zhao, X., Xu, J., et al. (2011). The role of runt-related transcription factor 2 (Runx2) in the late stage of odontoblast differentiation and dentin formation. Biochem Biophys Res Commun 410, 698–704.

41. Lin, C. R., Kioussi, C., O’Connell, S., Briata, P., Szeto, D., Liu, F., Izpisúa-Belmonte, J. C. and Rosenfeld, M. G. (1999). Pitx2 regulates lung asymmetry, cardiac positioning and pituitary and tooth morphogenesis. Nature 401, 279–282.

42. Liu, Z., Liu, S., Yao, J., Bao, L., Zhang, J., Li, Y., Jiang, C., Sun, L., Wang, R., Zhang, Y., et al. (2016). The channel catfish genome sequence provides insights into the evolution of scale formation in teleosts. Nat Commun 7, 11757.

43. MacDougall, M. (1998). Refined mapping of the human dentin sialophosphoprotein (DSPP) gene within the critical dentinogenesis imperfecta type II and dentin dysplasia type II loci. Eur J Oral Sci 106 Suppl 1, 227–233.

44. Magne, D., Bluteau, G., Lopez-Cazaux, S., Weiss, P., Pilet, P., Ritchie, H. H., Daculsi, G. and Guicheux, J. (2004). Development of an odontoblast in vitro model to study dentin mineralization. Connect Tissue Res 45, 101–108.

45. Mammoto, T., Mammoto, A., Torisawa, Y. S., Tat, T., Gibbs, A., Derda, R., Mannix, R., de Bruijn, M., Yung, C. W., Huh, D., et al. (2011). Mechanochemical control of mesenchymal condensation and embryonic tooth organ formation. Dev Cell 21, 758–769.

46. Martin, K. J., Rasch, L. J., Cooper, R. L., Metscher, B. D., Johanson, Z. and Fraser, G. J. (2016). Sox2+ progenitors in sharks link taste development with the evolution of regenerative teeth from denticles. Proc Natl Acad Sci U S A 113, 14769–14774.

47. Matalová, E., Lungová, V. and Sharpe, P. (2015). *Development of Tooth and Associated Structures*: Elsevier Inc.

48. Miyake, T., Vaglia, J. L., Taylor, L. H. and Hall, B. K. (1999). Development of dermal denticles in skates (Chondrichthyes, Batoidea): patterning and cellular differentiation. J Morphol 241, 61–81.

49. O’Neill, P., McCole, R. B. and Baker, C. V. (2007). A molecular analysis of neurogenic placode and cranial sensory ganglion development in the shark, Scyliorhinus canicula. Dev Biol 304, 156–181.

50. Rasch, L. J., Martin, K. J., Cooper, R. L., Metscher, B. D., Underwood, C. J. and Fraser, G. J. (2016). An ancient dental gene set governs development and continuous regeneration of teeth in sharks. Dev Biol 415, 347–370.

51. Reif, W.-E. (1982a). Evolution of dermal skeleton and dentition in vertebrates. Evolutionary biology 15, 287–368.

52. Reif, W. E. (1982b). Morphogenesis and function of the squamation in sharks. Neues Jahrbuch für Geologie und Paläontologie-Abhandlungen, 172-183.

53. Rivera-Rivera, C. J., Guevara-Delgadillo, N. I., Bahechar, I. A., Shea, C. A. and Montoya-Burgos, J. I. (2021). Loricarioid catfish evolved skin denticles that recapitulate teeth at the structural, developmental, and genetic levels. bioRxiv, 2021.2005.2017.444419.

54. Rivera-Rivera, C. J. and Montoya-Burgos, J. I. (2017). Trunk dental tissue evolved independently from underlying dermal bony plates but is associated with surface bones in living odontode-bearing catfish. Proc Biol Sci 284.

55. Sire, J. Y. and Allizard, F. (2001). A fourth teleost lineage possessing extra-oral teeth: the genus atherion (teleostei; atheriniformes). Eur J Morphol 39, 295–305.

56. Sire, J. Y. and Huysseune, A. (1996). Structure and development of the odontodes in an armoured catfish, Corydoras aeneus (Siluriformes, Callichthyidae). Acta Zoologica 77, 51–72.

57. Sire, J. Y., Marin, S. and Allizard, F. (1998). Comparison of teeth and dermal denticles (odontodes) in the teleost Denticeps clupeoides (Clupeomorpha). J Morphol 237, 237–255.

58. St Amand, T. R., Zhang, Y., Semina, E. V., Zhao, X., Hu, Y., Nguyen, L., Murray, J. C. and Chen, Y. (2000). Antagonistic signals between BMP4 and FGF8 define the expression of Pitx1 and Pitx2 in mouse tooth-forming anlage. Dev Biol 217, 323–332.

59. Thesleff, I. and Tummers, M. (2008). Tooth organogenesis and regeneration. In StemBook. Cambridge (MA): Harvard Stem Cell Institute.

60. Thompson, A. C., Capellini, T. D., Guenther, C. A., Chan, Y. F., Infante, C. R., Menke, D. B. and Kingsley, D. M. (2018). A novel enhancer near the Pitx1 gene influences development and evolution of pelvic appendages in vertebrates. Elife 7:e38555.

61. Tucker, A. S., Headon, D. J., Schneider, P., Ferguson, B. M., Overbeek, P., Tschopp, J. and Sharpe, P. T. (2000). Edar/Eda interactions regulate enamel knot formation in tooth morphogenesis. Development 127, 4691–4700.

62. Welten, M., Smith, M. M., Underwood, C. and Johanson, Z. (2015). Evolutionary origins and development of saw-teeth on the sawfish and sawshark rostrum (Elasmobranchii; Chondrichthyes). R Soc Open Sci 2, 150189.

63. Williamson, W. C. (1849). On the microscopic structure of the scales and dermal teeth of some ganoid and placoid fish. Philosophical Transactions of the Royal Society of London 139, 435–475. ---- (1851). Investigations into the structure and development of the scales and bones of fishes. In Abstracts of the Papers Communicated to the Royal Society of London, pp. 969-971: The Royal Society London.

64. Witten, P., Sire, J. Y. and Huysseune, A. (2014). Old, new and new-old concepts about the evolution of teeth. Journal of Applied Ichthyology 30, 636–642.

65. Xie, K. T., Wang, G., Thompson, A. C., Wucherpfennig, J. I., Reimchen, T. E., MacColl, A. D. C., Schluter, D., Bell, M. A., Vasquez, K. M. and Kingsley, D. M. (2019). DNA fragility in the parallel evolution of pelvic reduction in stickleback fish. Science 363, 81–84.

66. Yamakoshi, Y. (2008). Dentin Sialophophoprotein (DSPP) and Dentin. J Oral Biosci 50, 33–44.

